# Functioning human lung organoids model pulmonary tissue response from carbon nanomaterial exposures

**DOI:** 10.1101/2023.03.30.534957

**Authors:** Rahaf Issa, Neus Lozano, Kostas Kostarelos, Sandra Vranic

**Affiliations:** Nanomedicine Lab, Faculty of Biology, Medicine and Health, The University of Manchester, Manchester, M13 9PT, U.K; National Graphene Institute, The University of Manchester, Booth Street East, Manchester, M13 9PL, U.K; Catalan Institute of Nanoscience and Nanotechnology (ICN2), CSIC and BIST, Campus UAB, Bellaterra, 08193 Barcelona, Spain

**Keywords:** Lung organoid, Carbon nanomaterial (GO and MWCNT), Pulmonary exposure, Lung fibrosis, Alternative toxicology model, Nanotoxicity.

## Abstract

Human lung organoids (HLOs) are increasingly used to model development and infectious diseases, however their ability to recapitulate functional pulmonary tissue response to nanomaterial (NM) exposures has yet to be demonstrated. Here, we established a lung organoid exposure model that utilises microinjection to present NMs into the lumen of organoids. Our model assures efficient, reproducible and controllable exposure of the apical pulmonary epithelium, emulating real-life human exposure scenario. By comparing the impact of two well studied carbon-based NMs, graphene oxide sheets (GO) and multi-walled carbon nanotubes (MWCNT), we validated lung organoids as tools for predicting pulmonary NM-driven responses. In agreement with established *in vivo* data, we demonstrate that MWCNT, but not GO, elicit adverse effects on lung organoids, leading to a pro-fibrotic phenotype. Our findings reveal the capacity and suitability of HLOs for hazard assessment of NMs, aligned with the much sought-out 3Rs (animal research replacement, reduction, refinement) framework.

## 1.0 Introduction

The current standard for pulmonary tissue response analysis combines mono/co-culture *in vitro* systems^[1,2]^, with time-intensive and ethically charged *in vivo* models^[3–5]^. Though mono/co-culture systems allow for large-scale screening of nanomaterials (NMs), and have provided a better understanding of cellular effects with regards to physicochemical features of NMs^[2]^, such systems are often too simplistic. They lack physiological functions and only partially depict the complex cross-talk between cells. Such models also do not represent the complexity of human pulmonary epithelium and may overestimate the toxic potential of NMs. With *in vivo* models, ethical concerns and costs limit the number of NMs that can be evaluated. Furthermore, concerns regarding animal welfare, suitability and the accuracy of such models to predict human physiological responses have initiated much discussion around the use of animals in chemical safety testing, and the application of the 3Rs framework which aims to reduce, refine and replace animal testing^[6]^.

In recent years, advanced three-dimensional *in vitro* cultures have grown in popularity^[7]^. Examples include *ex vivo* precision-cut lung slices, lung cell type spheroids, air-liquid interface cultures and microfluid lung-on-a-chip models^[7]^. Such platforms have provided a unique alternative to bridge the gap between traditional two-dimensional *in vitro* and *in vivo* models, but to date most consist of only two or three cell types^[8–10]^. It is therefore essential to develop a new approach that can accurately model NM-induced pulmonary tissue response in relatively short-term^[11]^, providing sufficient cellular complexity and realistic NM-exposure scenario.

One of the most promising and emerging modelling technologies exploits the use of human organoid systems. Lung organoids are *in vitro,* self-organising, three-dimensional culture systems that are derived from either progenitor cells of the adult lung^[12]^ or pluripotent stem cells^[13,14]^. They recapitulate key structures and functions of human tissue, making it possible to study developmental patterns and diseases^[12,15]^. To date, several groups have described the step-wise differentiation of human pluripotent stem cells into lung organoids containing proximal and/or distal airway cells^[13,14,16–20]^. In terms of application, lung organoids have been used to better understand various pulmonary diseases, from cystic fibrosis to lung cancer^[12]^, and infectious diseases such as respiratory syncytial virus infection^[14]^ and SARS-CoV-2 infection^[21]^. Several perspective articles have also discussed the potential usefulness of human organoids in toxicology and hazard assessments^[22,23]^, with a focus on drug-induced organ toxicity in liver, heart, kidney, intestinal and brain organoids^[22]^. However, the ability of multilineage, multicellular HLOs to recapitulate functional pulmonary tissue response to NMs has not yet been demonstrated.

With the expansion of nanotechnology, there is a pressing need to better understand how NMs may impact human health. Inhalation of aerosolised materials is one of the first routes of exposure, with the respiratory system first impacted. Previous *in vitro* and *in vivo* studies have shown that some carbon nanotubes (CNTs) induce pulmonary-associated toxicity^[9,24–28]^, with their ability to biopersist in the lungs strongly influenced by their physicochemical properties (length, diameter, aspect ratio and rigidity)^[29]^. Long multi-walled CNTs (MWCNT), for example, have been shown to induce adverse pulmonary effects, causing persistent inflammation^[30]^, abnormal deposition of extracellular matrix^[31]^, formation of irreversible fibrotic lesions with tissue remodelling^[26]^, necrosis, and even carcinogenesis^[32]^.

As a safer alternative graphene oxide (GO), the oxidised form of graphene, has emerged^[33]^. GO has fast become one of the most studied two-dimensional NMs, mainly due to its ease of production, high surface area exploitable for drug loading, and oxygen-rich functional groups that enable its good colloidal dispersion in physiological *milieu*. Because of its similarity in chemical composition to other carbon nanostructures, namely CNTs, efforts are made to evaluate the impact of GO on lungs after inhalation^[4,5,34,35]^. Our group and others have previously shown that GO-induced toxicity is strongly dependent on the lateral dimension of the material^[1,2,36]^. Micrometric-sized GO sheets, when presented to rodent lung by a single intranasal aspiration, trigger granuloma formation that persists for up to 90d, but this does not lead to pulmonary fibrosis^[37]^. In contrast, nanometric-sized GO sheets are not associated with any adverse pulmonary effects^[37,38]^. More recently, it was shown that murine lungs can rapidly recover from repeated low dose exposures to micrometric GO sheets, despite initial signs of material-induced lung inflammation and genotoxicity^[4,5]^.

The potential adverse impact of NMs emphasises the need for improved hazard assessment testing platforms and models that are relevant to human physiology. In this study, we used HLOs to evaluate NM-induced pulmonary responses, and assess their suitability as a predicative tool in hazard assessment of carbon NMs. We adapted previously published protocols to develop an embryonic stem cell-derived HLO, composed of six major proximal and distal airway epithelial cells and mesenchymal cells which are important for modelling pulmonary fibrosis^[15]^. We established a novel NM microinjection protocol that tailored the dosing of carbon-based NMs to the apical surface of lung organoids, hence mimicking human pulmonary exposures. The impact of small (30-750nm, sGO) and large (11-67µm, lGO) GO sheets in lateral dimension was compared with long and rigid MWCNT on HLOs after 1d and 7d exposure. Key biological endpoints relevant to pulmonary tissue response of carbon-based NMs^[4,5,37]^ were investigated, including cytotoxicity, material-cell interactions, tissue remodelling and pulmonary fibrosis.

## 2.0 Materials and Methods

### 2.1 hESC maintenance

Man-5 (Manchester University Embryonic Stem Cell Line 5, RRID:CVCL_L193) were kindly provided by Professor Susan Kimber, University of Manchester. Cells were cultured on growth factor-reduced Matrigel (Corning), in mTeSR1 medium (STEMCELL Technologies). Media was changed daily and cells were passaged every 5-6 days using 0.5 mM EDTA. Media was supplemented with 10 µM ROCK inhibitor Y-27632 (Tocris) for a maximum of 24 h after splitting. Cells were maintained in an undifferentiated state at 37°C, 5% CO_2_, and used between passage 19-28. Cells were tested for mycoplasma contamination every 4 months. Karyotype was verified using Cell Guidance Systems Genetics Service on fixed cells of P28 Man-5.

### 2.2 Human lung organoid culture

Protocol for generation of lung organoids was adapted from Carvalho *et al.*^[18,39]^. All steps were carried out in serum-free differentiation (SFD) medium consisting of IMDM/F12 (3:1) (Life Technologies), N2 and B27 supplements (Life Technologies), 1% GlutaMax (Gibco), 1% penicillin-streptomycin (Sigma), 0.05% bovine serum albumin (Gibco), with 50 µg/ml ascorbic acid (Sigma) and 0.4 µM monothioglycerol (Sigma) added fresh. All growth factors were purchased from R&D systems, unless otherwise stated. Cultures were maintained at 37°C, 5% CO_2_. For induction of definitive endoderm, hESC (3-10 cells/clump) were plated onto low-attachment 6-well plates (Corning) in SFD medium supplemented with 10 µM Y-27632, 100 ng/ml activin A, 0.5 ng/ml BMP4 and 2.5 ng/ml FGF2. Cells were cultured for 84 h and media was changed daily. Endoderm yield was determined by the expression of CXCR4 and c-KIT, with cells used in all studies exhibiting >95% endodermal yield. On day 4.5, embryoid bodies were dissociated into single cells using 0.05% trypsin/0.02% EDTA and plated onto fibronectin-coated plates (∼60,000 cells/cm^2^). For induction of anterior foregut endoderm, cells were cultured in SFD medium supplemented with 10 µM SB431542 and 2 µM dorsomorphin dihydrochloride for 24 h, then switched to 10 µM SB431542 and 1 µM IWP2 for a further 24 h. For induction of early-stage lung progenitor cells (day 6-15), cells were exposed to SFD medium supplemented with 3 µM CHIR99021, 10 ng/ml FGF7, 10 ng/ml FGF10, 10 ng/ml BMP4 and 50 nM all-trans retinoic acid (Sigma). Media was changed every other day. On day 15, the early lung progenitors were briefly trypsinised, and cell clumps (1-2 mm) were replated onto growth factor-reduced Matrigel-coated plates at a 1:2 split ratio. Cells were cultured in SFD medium supplemented with 3 µM CHIR99021, 10 ng/ml FGF7 and 10 ng/ml FGF10 until day 25, with media changes every other day. For differentiation of lung progenitors into organoids (day 25-50), cells were briefly trypsinised and embedded in 4.5 mg/ml rat tail collagen I (Corning, Col I). Gels were allowed to set for 15 min, before SFD medium supplemented with 10 ng/ml FGF7, 10 ng/ml FGF10, 50 ng/ml dexamethasone, 0.1 mM 8-bromo-cAMP and 0.1 mM IBMX was added. Media was changed every other day.

### 2.3 Material preparation

Aqueous suspensions of small (s) and large (l) GO sheets were produced in house as previously described^[3]^. Detailed characterisation of sGO and lGO can be found in **Figure S3**. Optical image was acquired with a Nikon Eclipse LV100 microscope in transmittance mode at a magnification of 50x, at the ICN2 Molecular Spectroscopy and Optical Microscopy Facility. Electron microscopy images were obtained using a Magellan 400L field emission scanning electron microscope (Oxford Instruments), at the ICN2 Electron Microscopy Unit, with an Everhart–Thornley secondary electrons detector, using an acceleration voltage of 20 kV and a beam current of 0.1 nA. Atomic force microscopy height image was taken using an Agilent 5500 AFM/SPM microscope at the ICN2 Scanning Probe Microscopy Facility, in tapping mode equipped with silicon cantilevers (Ted Pella) with a nominal force of 40 N m−1 and a resonance frequency of 300 kHz. Multi-walled carbon nanotubes (Mitsui-7, MWCNT) were a kind gift from Professor Ulla B Vogel, National Research Centre for the Working Environment, Denmark. MWCNT were exposed to dry heat sterilisation (160°C for at least 16 h) to inactivate any contaminating endotoxins. MWCNT stock suspension was prepared in phosphate buffer saline without Ca^2+^ and Mg^2+^ (Sigma, PBS^-/-^) containing 0.5% BSA and subjected to sonication for 5-7 min at nominal 80 W. GO and MWCNT suspensions were diluted in PBS^-/-^ and used immediately. All nanomaterials were evaluated for endotoxin contamination as described in Mukherjee *et al.*^[40]^ and tested negative (data not shown).

### 2.4 Microinjection of NMs

Microcapillaries (3.5”, Drummond) were prepared using the P-97 micropipette puller (Sutter) fitted with a 3.0 mm trough filament; with pressure set at 300, heat at 295, velocity at 10 and delay at 250. Pulled capillaries were UV-sterilised for 20 min before use. Only microneedles with a tip diameter between 8-21 µm were used in the study. Prior to injections, organoids were gently transferred to an 8-well chamber slide (Ibidi) using a wide-bore pipette tip (2-3 organoids/well). Fresh 4.5 mg/ml rat tail Col I was added to each well and allowed to gel to prevent movement during injection. Organoids were then imaged using the EVOS FL imaging system (Thermo Scientific) at 4x magnification. Organoid area was measured using the image analysis software, FiJi (v2.9.0/1.53t). Images were converted to 8-bit and threshold values were determined to cover the organoid surface area. The area (mm^2^) was determined using the ‘Analyse Particles’ function of FiJi. For studies assessing the organoid epithelial barrier, 4kDa FITC-dextran (FD4, Sigma) was prepared at 2 mg/ml in PBS and injected into the organoids at a volume of up to 400 nl/mm^2^. Treatments (sGO, lGO, MWCNT or PBS^-/-^ vehicle control) were injected into the organoids, at a maximum volume of 100 nl/mm^2^, using the Nanoject II microinjector (Drummond). To inject the airspace-like lumen of the organoids, we used a diagonal needle approach. The microneedle was first lowered on the organoid using a micromanipulator, and pressed down onto the organoid surface. The needle tip was easily observed in the focal plane of the target organoid. The microneedle was then moved laterally to pierce the organoid. The contents of the loaded microneedle were injected into the organoid lumen. The microneedle was finally removed from the organoid lumen, by moving up and laterally away. Successful injections into the luminal space were confirmed by the slight expansion of the organoid upon injection. A minimum of 4 independent replicates, each containing 3 organoids were used.

### 2.5 Quantification of cell number per organoid

Lung organoids in Col I gels were digested with 150 U/ml collagenase type I (Gibco, in IMDM) for 1 h at 37°C. Organoids were collected and further dissociated with pre-warmed 0.05% trypsin/EDTA and gentle pipetting into single cells. Cells were then counted using a haemocytometer. A total of 31 organoids, ranging between 0.2 to 3.5 mm^2^, were analysed.

### 2.6 Histology

Col I gels containing the lung organoid were embedded in Optical Cutting Temperature (OCT) and snap-frozen in isopentane (Sigma) over dry ice. Serial sections were obtained at 10 µm thick using a cryostat (Leica CM1950). Sections were either stained with haematoxylin and eosin, using the XL autostainer (Leica Biosystems), or stored at -80°C for immunostaining later. Brightfield images were acquired on a 3D-Histech Pannoramic-250 microscope slide scanner using a 20x/0.80 Plan Apochromat objective (Zeiss). Snapshots of the slide-scans were processed using the Case Viewer software (3D-Histech).

### 2.7 Immunofluorescence

Sections were briefly fixed with 95% ethanol for 1 min followed by 4% paraformaldehyde (Thermo Scientific) for 15 min. After washing twice with PBS, sections were permeabilised with 0.25% triton X-100 in blocking buffer (5% goat or donkey serum in PBS^-/-^) for 10 min, and then incubated with blocking buffer for 1 h. Primary antibodies (Table S1) diluted in blocking buffer were incubated at 4°C overnight in a humidified chamber. The following day, sections were washed with PBS^-/-^ 3×5 min, incubated with the secondary antibodies (Table S2, prepared in blocking buffer) for 2 h at room temperature followed by 10 min incubation with DAPI. Slides were again washed with PBS^-/-^ 3×5 min and mounted using Prolong Gold Antifade Mountant (Thermo Scientific). Sections stained with secondary only antibodies were included to determine fluorescence threshold before image acquisition in each channel. Images were acquired using Olympus BX63 upright microscope (20x 0.75 UApo/340 objective) and captured using a DP80 camera through CellSens Dimension software (Olympus). Specific band pass filter sets for DAPI, FITC, Texas red and Cy5 were used to prevent spectral bleed-through from one channel to the next.

For 2D cultures (hESCs or d25 lung progenitors), cells were fixed with only 4% paraformaldehyde for 15 min prior to permeabilisation and staining. Images were acquired using Zeiss LSM 780 confocal microscope (10x 0.50 Flar objective or 20x 0.8 Plan-Apochromat objective, 1.5x confocal zoom).

For assessing cell-material interactions, organoid sections were stained for markers of proximal and distal airway cells after exposure to s/lGO and MWCNT. Sequential organoid sections, 10 µm apart, were stained for the following cells types and markers; multiciliated (acTUB), goblet (MUC5AC), club (SCGB1A1), basal (KRT5), AECI (HOPX), AECII (MUC1), secretory mucins (MUC5AC/MUC1) and surfactant (SFTPB). Brightfield with immunofluorescence images were overlayed to confirm the localisation of the materials, as they appear black/brown under optical light.

### 2.8 Quantification of immunofluorescence

Images of each marker were quantified using the FiJi software (v2.9.0/1.53t). Images were converted to 8-bit and the threshold was adjusted to correspond to the nuclear stain, which allows for measurement of total area. For cytoplasmic or membrane markers, the threshold was adjusted to cover the stained area. Fluorescent area was analysed by the ‘Analyse Particles’ function of FiJi. Cells staining positive for each marker were calculated by dividing the total area of positive cells over the total area of DAPI.

### 2.9 Flow cytometry

Cell colonies were dissociated with either StemPro accutase (Gibco) for 8 min (hESC) or TrypLE (Gibco) for 15 min (d15/25 lung progenitors) at 37°C. Cells were collected and washed twice with FACS buffer (0.5% BSA and 2 mM EDTA in PBS^-/-^). For pluripotency assessment of extracellular markers, cells were incubated with unconjugated anti-SSEA4 or anti-TRA-1-60 for 1 h on ice, followed by incubation with goat anti-mouse Alexa Fluor 488 for 30 min on ice. For intracellular markers (Oct.4, Nanog and SOX2), cells were fixed with 2% paraformaldehyde for 20 min and permeabilised with 0.1% triton X-100 (in PBS^-/-^) for 15 min before staining. Donkey anti-rabbit Alexa Fluor 488 (for Oct.4) and goat anti-mouse Alexa Fluor 488 (for SOX2 and Nanog) were used to label the cells. Cells stained with secondary only antibodies were also included for gating. For analysis of d15/25 lung progenitors, cells were incubated with anti-EpCAM Alexa Fluor 488, anti-PDPN PE and anti-MUC1 APC at 4°C for 45 min. Details for antibodies and dilutions used can be found in Table S3. Cells were analysed on BD Fortessa or BD Fortessa X20 (BD Bioscience). Results were analysed using Flowjo v10.2 software. Analysis was gated on live, doublet-excluded cells.

### 2.10 Annexin V/propidium iodide staining

After material exposure and digestion of Col I gels, organoids were dissociated with pre-warmed 0.05% trypsin/EDTA and gentle pipetting into single cells. The cell suspension was incubated with 10% FCS (Sigma) for 30 min to allow the cell membrane to recover from trypsinisation and prevent false positive staining with Annexin V (AV). Cells were then washed with ice-cold PBS. AV staining was performed according to the manufacturer (Molecular Probes). Briefly, cells (Σ1×10^5^) were resuspended in 100 µl AV binding buffer and stained with 3 µl AV-Alexa Fluor 488 conjugate for 15 min. Propidium iodide (PI, Sigma) was added immediately before analysis at a final concentration of 1.5 µg/ml. Cells were analysed on BD Fortessa X20 (BD Bioscience). Results were analysed using Flowjo v10.2 software. Analysis was gated on doublet-excluded cells (gating strategy can be found in **Figure S6A**).

### 2.11 Live imaging

Lung organoids in Col I gels were digested with 150 U/ml collagenase type I (in IMDM) for 1 h at 37°C. Organoids were collected and further dissociated with pre-warmed 0.05% trypsin/EDTA and gentle pipetting to small sized cell clumps. Cell clumps were resuspended in SFD medium containing 10 ng/ml FGF7, 10 ng/ml FGF10, 50 ng/ml dexamethasone, 0.1 mM 8-bromo-cAMP and 0.1 mM IBMX, and plated in a CELLview culture dish (Greiner). Live imaging was performed using an Olympus IX83 inverted microscope (60x 0.70LUC PlanFL N UIS 2 (Ph2) objective) equipped with an Orca ER camera (Hamamatsu) through CellSens software (Olympus).

### 2.12 Transmission electron microscopy

Col I gel-free organoids were fixed with 4% PFA and 2.5% glutaraldehyde in 0.1 M HEPES buffer (pH 7.2) for at least 2 h, and post-fixed with ferrocyanide reduced osmium. Organoids were dehydrated in increasing concentrations of ethanol (30-100%) and in 100% acetone, before embedding in TAAB LV resin and polymerisation at 60°C (>24 h). Ultrathin sections (70 nm) were cut with a diamond knife (Diatome) and mounted on copper grids. Organoids were examined using a Talos L120C electron microscope (Thermo Fisher).

### 2.13 Statistical analysis

Data were analysed using GraphPad Prism 9 (GraphPad Software Inc). For multiple group comparisons, one- or two-way ANOVA was performed followed by Dunnett or Tukey tests. Where appropriated, unpaired two-tailed Student’s t-test was performed. *P*<0.05 was considered statistically significant.

## 3.0 Results

### 3.1 HLOs development and characterization

A multilineage and mature HLO was established by adapting previously published protocols (**Figure 1A**)^[18,39]^. The hESC line used in this study, Man-5 (**Figure 1B**), was maintained in a pluripotent state and exhibited a normal female karyotype **(Figure S1)**. Differentiation of cells was verified at key stages; d4.5, d15 and d25. At d4.5, >95% cells were CXCR4^+^cKIT^+^, markers of definitive endoderm (**Figure 1C**). By d15, 60% cells were epithelial (EpCAM^+^) and co-expressed pre-AECII markers, MUC1^+^PDPN^+^ (**Figure 1D**). The purity of the lung progenitor cell population further increased and at d25, >90% cells were EpCAM^+^ with 28% expressing early AECII markers, MUC1^+^PDPN^-^ (**Figure 1E**). The protocol also yielded cultures that are highly enriched for the expression of the endodermal marker FOXA2 and lung transcription factor NKX2.1, as well as transcription factors SOX2 and SOX9 which are involved in proximodistal patterning of the lung (**Figure 1F**).

**Figure 1.**
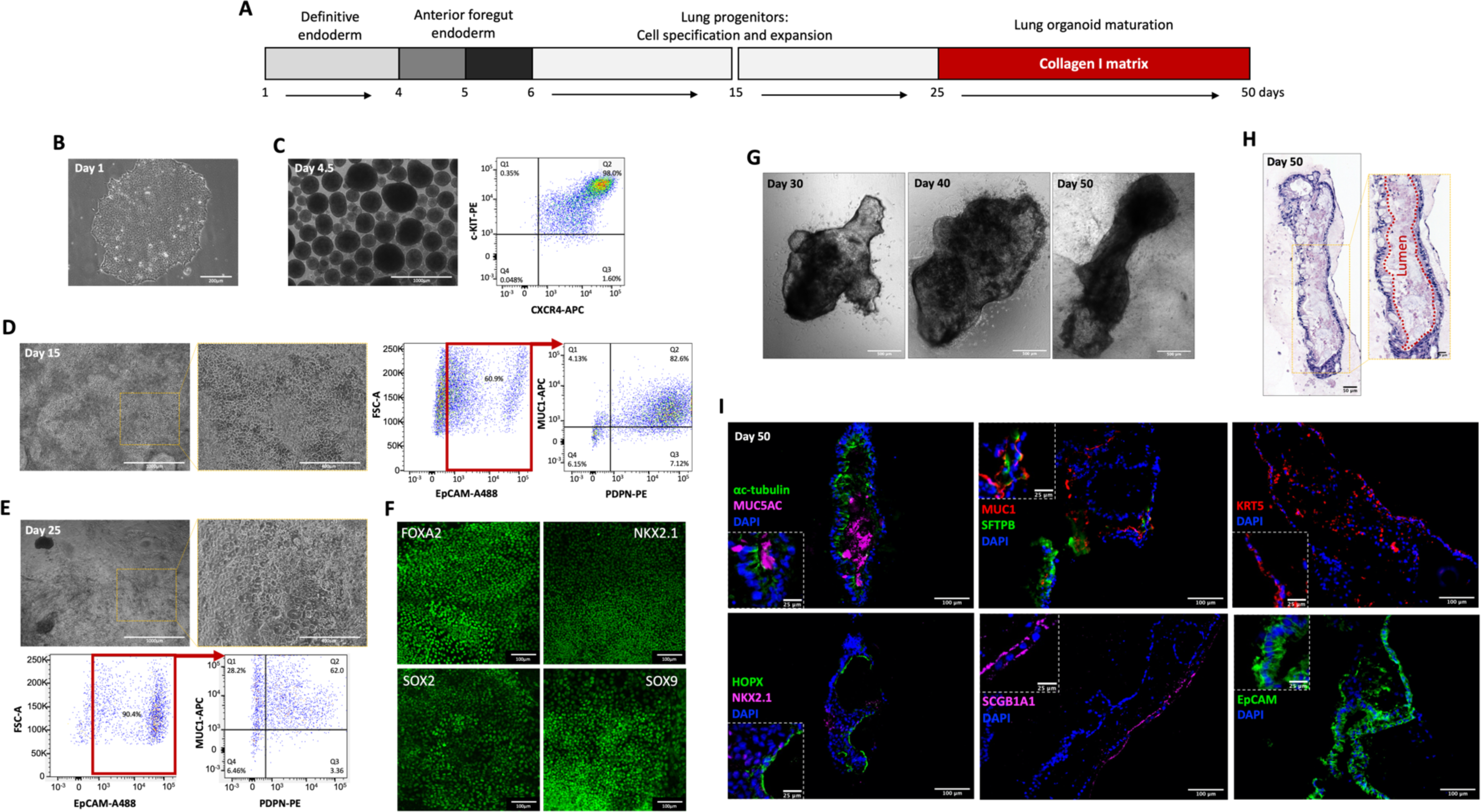
Generation of multilineage lung organoids from human embryonic stem cells (hESCs) **A**, Schematic representation of the differentiation protocol, adapted from Carvahlo *et al*.^[39]^, from day 1 to 50. **B,** Brightfield image of hESC line Man5 after culturing to confluence. Scale bar, 200µm. **C,** Brightfield image of definitive endoderm at day 4.5, and a representative example of the flow cytometry profile after staining at day 4.5 cells for markers of definitive endoderm, CXCR4 and cKIT. Scale bar, 1000µm. **D-E**, Brightfield images of day 15 **(D)** and day 25 **(E)** of lung progenitors. Cell population profile after staining at day 15 (**D**) and day 25 (**E**) lung progenitor cells for EpCAM, and markers of pre- or mature-alveolar cells, PDPN and MUC1, respectively. Scale bar, 1000µm (Inset, 400µm). **F**, Immunofluorescence micrographs of day 25 lung progenitors expressing markers of lung lineage, FOXA2, NKX2.1, SOX2 and SOX9. Scale bar, 100µm. **G**, Brightfield image of developing lung organoids at day 30, 40 and 50. Scale bar, 500µm. **H**, H&E staining of a day 50 lung organoid (cryo-sectioned, 6µm slices). Scale bar, 50µm (Inset, 20µm). **I**, Immunofluorescence micrographs of day 50 lung organoid sections expressing markers of proximal and distal airway epithelial cells. Multiciliated cells (ac-tubulin), goblet (MUC5AC), club (SCGBA1A), basal (KRT5), AECI (HOPX), AECII (MUC1, SFTPB), and lung epithelium (EpCAM). Scale bar, 100µm (Inset, 25µm). FSC-A, forward scatter area. A488, Alexa-fluor 488. PE, phycoerythrin. APC, allophycocyanin. **Movie 1. Live imaging of multi-ciliated cells at d52 after collagenase I digestion and brief trypsinisation.** Video shows a looped 10s snippet. Scale bar, 25µm.

After maturation in a 3D collagen matrix for 50d (**Figure 1G**), HLOs exhibited an airspace-like lumen void of cells (**Figure 1H**), that may be exploited for controlled NM exposures. Cells within the organoid were characterised and the organoid functional features were assessed. Functional ciliated cells, which can be seen beating when the organoid is dissociated into small cell clumps were identified (**Movie 1**). Using immunofluorescence staining and TEM, the six major proximal and distal pulmonary epithelial cell types were detected (**Figures 1I, S2**). Basal (KRT5^+^) and club (SCGB1A1^+^) cells were interspaced throughout the organoid, whilst ciliated (acTUB^+^) and goblet cells (MUC5AC^+^) appeared in clusters (with secreted mucin accumulating in the luminal space). Closer examination revealed that markers occurred in epithelial structures that appeared polarised, with KRT5^+^ cells on the basal side and acTUB^+^ on the apical side. Distal AECII cells expressing SFTPB, SFTPC and MUC1 were abundant in the culture. Markers of AECI such as HOPX were found in areas of cell flattening and thinning of the cultures and exhibited thin cytoplasmic extensions, which is a morphological characteristic of AECI *in vivo* cells.

Consistent with the immunofluorescence data, TEM showed the presence of goblet cells with secretory granules and ciliated cells reaching the lumen **(Figure S2)**. Cells with electron-dense lamellar bodies, the functional organelles in which surfactant is stored before exocytosis into the air spaces, were evident and indicative of ACEII maturation. In some areas, surfactant deposition was also observed **(Figure S2)**. Taken together these findings confirm the presence of functional proximal and distal airway cells in the lung organoids.

### 3.2 Lung organoid exposure model

In lung organoids, the apical surface of the epithelium is enclosed while the basolateral surface faces the outside of the organoid (**Figure 1H**). We established a protocol to microinject a controlled volume and dose of NMs into the lumen of the organoid, simulating a real pulmonary exposure scenario of the apical region of the lung epithelium (**Figure 2A**). To control the amount of NM injected per organoid, we considered the following parameters: organoid size, cell number per organoid, volume injected, NM dose, and vehicle. We first addressed the variation in organoid size in relation to the cell number per organoid, as organoid development is hindered by the lack of size uniformity. Using FiJi image analysis software, the size of 31 organoids was determined and ranged from 0.2 to 3.5 mm^2^ (**Figure 2B**). An excellent correlation between organoid size (measured by FiJi) and the actual number of cells in each organoid (after dissociation into single cells) was observed, with R^2^ value of 0.8654 (**Figure 2Bv**). To assess if microinjection compromises the epithelial barrier of the organoid, the retention of FD4 which is impermeable to intact membrane and intracellular junctions was assessed (**Figure 2C**). This also allowed us to evaluate whether the injected cargo remains in the organoid lumen after needle withdrawal. Injections of up to 100 nl/mm^2^ FD4 were retained by the organoids. However, increasing the volume to ≥200 nl/mm^2^ resulted in a compromise in epithelial barrier function, as evidenced by leaching of the fluorescent signal into surrounding matrix (**Figure 2C**).

**Figure 2.**
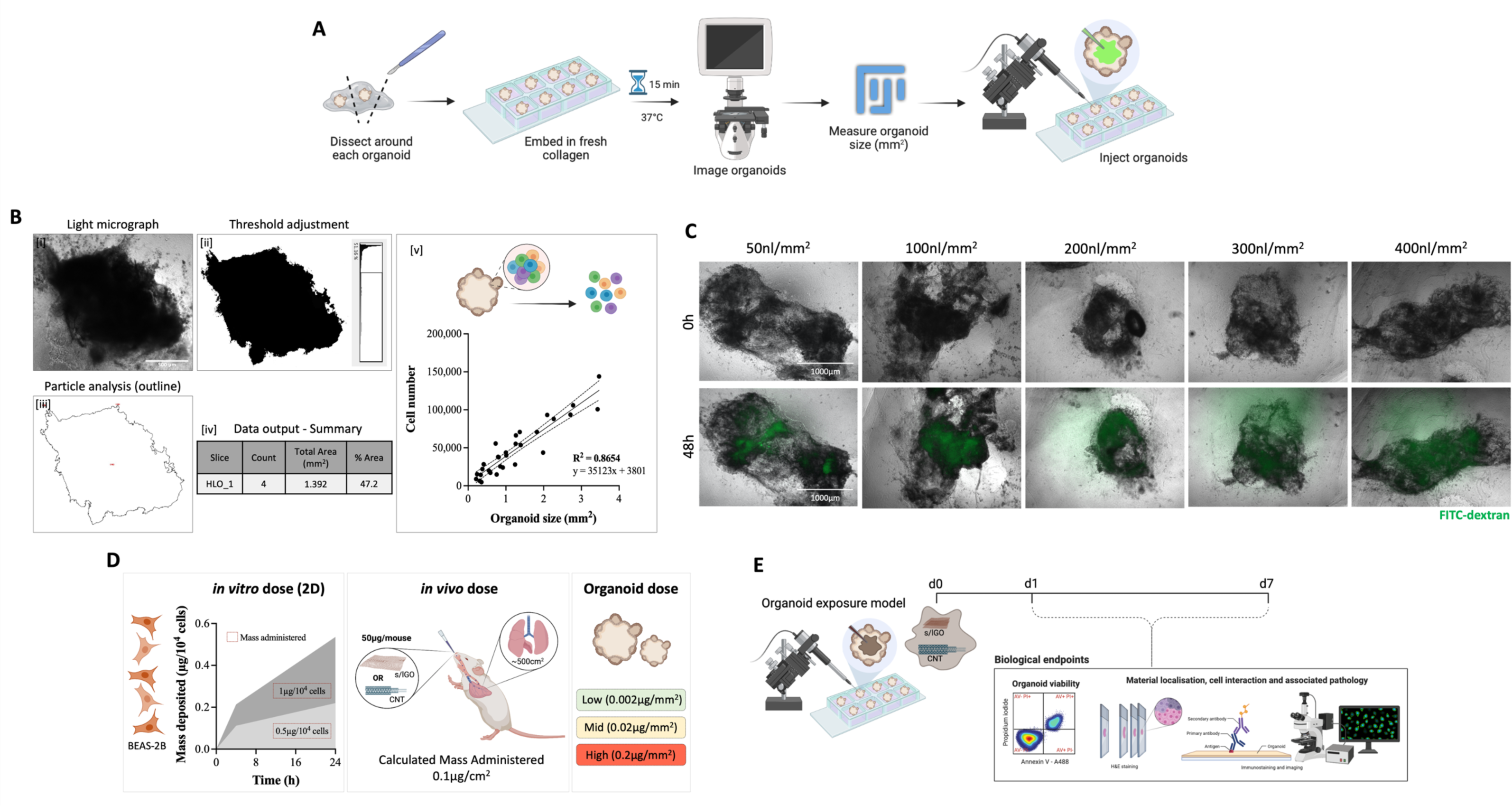
Microinjection of human lung organoids. **A,** Experimental set-up for cargo delivery into lung organoids. **B,** Correlation between organoid size (measured using the FiJi image analysis software [i-iv]) and total cell number [v], after organoids are dissociated into single cells. **B[i],** Light micrograph of a lung organoid. **B[ii],** After threshold adjustment in FiJi. **B[iii],** Organoid outline generated using the ‘Analyse Particles’ function in FiJi. **B[iv],** Summary of the data output detailing organoid size (mm^2^). Scale bar, 500 µm. **C,** Microinjection of varying volumes of FITC-dextran in lung organoids, before (0h) and after (48h) injection. Both panels show events under the GFP filter merged with brightfield. n = 2 organoids per condition. Scale bar, 1000 µm. **D,** Mass deposition reported *in vitro* for sGO on human lung epithelial cells, BEAS-2B, after 24h of exposure^[2]^, *in vivo* for s/lGO and MWCNT in an adult C57BL/6 mouse model after intranasal instillation^[37]^, and the concentrations of GO injected into human lung organoids. **E,** Schematic representation of the study design. Organoids were microinjected with sGO, lGO or MWCNT and exposed to the materials for 1d and 7d. The impact of material exposure on lung organoids was assessed by investigating key biological endpoints relevant to pulmonary toxicity of carbon-based NMs, which included cytotoxicity, material-cell interaction, tissue remodelling and pulmonary fibrosis.

To determine the NM dose to deliver into the organoids, we considered the amount of material that deposits per cell *in vitro*^[2]^, and the amount of material administered in previous *in vivo* studies^[37]^ (**Figure 2D**). Using UV-VIS spectrometry, the deposition of sGO in human bronchial epithelial cells, BEAS-2B, was calculated at ∼0.22 µg/10^4^ cells after 24h incubation (see Figure S2 in Chen *et al.*^[2]^). The dose *in vivo* was also estimated to be ∼0.1 µg/cm^2^, by taking into account the entire surface area of the respiratory tract of adult mouse (∼500 cm^2^) and the amount of material administered (50 µg/mouse) in previous *in vivo* studies using the same materials^[37]^. Thus, in the lung organoid exposure model here that contains 40,000 cells/mm^2^, a high GO dose equated to 0.2 µg/mm^2^, middle to 0.02 µg/mm^2^ and low to 0.002 µg/mm^2^. The volume injected (nl) was also adjusted based on the calculated organoid size (mm^2^) to prevent damage to the organoid’s epithelial barrier, with maximum volumes set to 100 nl/mm^2^.

The NMs used were thoroughly characterised in water and in the vehicle used for microinjections **(Figures S3-S4)**. PBS^-/-^ was selected as a suitable vehicle for the NMs. Both sGO and lGO did not show any major signs of agglomeration in PBS^-/-^, even after 24h of preparation **(Figure S4)**. By considering the following parameters: injection volume, organoid size, cell number per organoid, NM dose and vehicle used – we could tailor NM delivery to each organoid. For example, a 0.5 mm^2^ organoid exposed to high dose sGO received 0.1 µg sGO in 50 nl PBS^-/-^, whereas a 3 mm^2^ received 0.6 µg sGO in 300 nl PBS^-/-^.

A detailed schematic representation of the study design is depicted in **Figure 2E**.

### 3.3 s/lGO and MWCNT exposure does not permanently impact organoid viability

Cytotoxicity is a key endpoint when evaluating the toxicological profile of NMs. To assess the impact of injection and material exposure on organoid viability, we dissociated the organoids into single cells after s/lGO, MWCNT or PBS exposure and analysed the cells using the AV/PI flow cytometry assay. We included organoids exposed to PBS-alone to determine if puncture with the microinjection needles induces any mechanical trauma that may impact organoid viability. We also included organoids exposed to 2% Triton X-100 as a positive control for cell death. All three materials induced a short-lived apoptotic response that did not progress to late apoptosis or necrosis **(Figures S5A-C)**. At the highest dose tested for s/lGO and MWCNT, ∼20% more cells stained positive for markers of early apoptosis (AV^+^PI^-^) after 1d exposure when compared to the non-injected control. Furthermore, there was limited/no evidence of late apoptotic (AV^+^PI^+^) or necrotic (AV^-^PI^+^) cells after 1d material exposure. We did not observe a dose-dependent effect, suggesting that material-induced toxicity at 1d exposure may be limited to a few cell types within the organoids. The effects observed were not due to mechanical trauma as there was no significant difference between the non-injected and PBS-injected controls after both 1 and 7d exposure. Interestingly, after 7d exposure there was no evidence of any cell death (apoptosis, late apoptosis, or necrosis) in all material exposure groups. Taken together, these results suggest that lung organoids can withstand material exposure and have the capability to recover from the initial impact of NM exposure.

### 3.4 MWCNT but not GO cause remodelling of the organoid epithelium

Some carbon-based NMs, particularly MWCNT, are known to induce histopathological features *in vivo* from day 7 of exposure^[37]^. In organoids exposed to MWCNT, and not s/lGO, a significant reaction by the organoid parenchyma was observed (**Figures 3, S7-S8)**. Cells within MWCNT-exposed organoids were more densely packed with a distinctly thicker epithelium. Furthermore, the matrix appeared more disorganised, particularly after 7d exposure (**Figures 3B, S7B, S8B)**. In areas of high material load, there was significant damage to the epithelium as evidenced by the lack of cellular integrity (**Figure 3B**). Such organoids (MWCNT, 0.1 µg/mm^2^) were challenging to cryosection, as the nanotubes when present in high concentrations caused tissue damage (dragging and tearing) on sectioning. To reduce mechanical-related damage, only organoids exposed to Σ0.01 µg/mm^2^ MWCNT were subsequently used.

**Figure 3.**
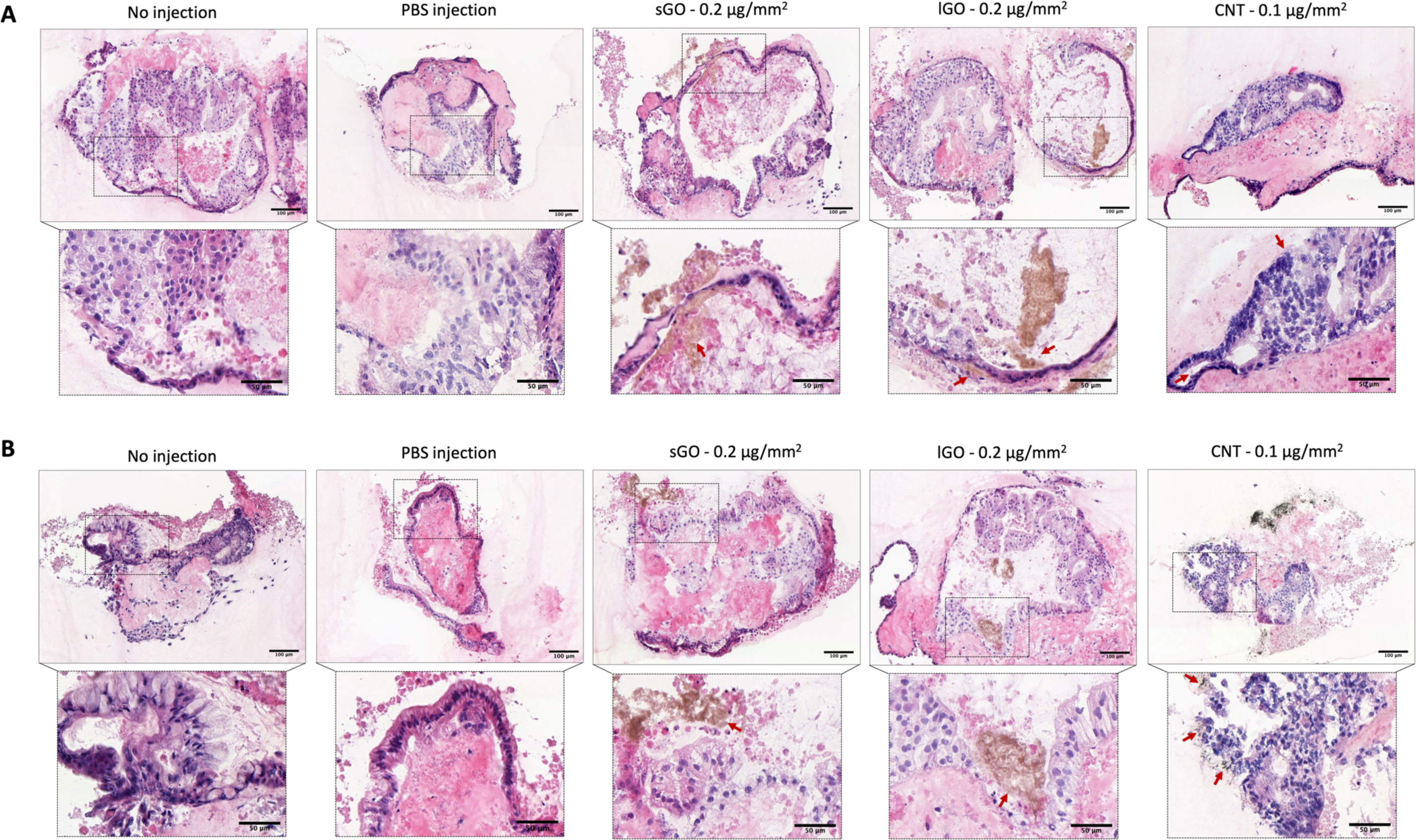
Histological analysis of human lung organoids after 1d and 7d exposure to high dose of carbon-based NM. **A,** 1d exposure. **B,** 7d exposure. Organoids were flash-frozen, cryo-sectioned at a thickness of 10 µm and stained with H&E. Arrows indicate material localisation. Scale bar, 100 µm (Inset, 50 µm). All three NMs were easily located in various regions of the organoids, appearing as brown (for GO) and black (for MWCNT) regions associated with the cells, extracellular matrix or secretory proteins in the luminal space. Histological analysis of lung organoids exposed to low- and mid-dose carbon-based NM are shown in Figure S7 and S8, respectively.

### 3.5 GO sheets interact less than MWCNT with alveolar cells

The interaction of NMs with cells can induce cellular responses that define the biological fate and toxicological profile of the NM^[1,2]^. To assess cell-material interactions, we stained the organoid sections for markers of proximal and distal airway cells after exposure to sGO, lGO and MWCNT.

Interestingly, all three materials interacted with only distal airway cells, AECI and ACEII (**Figures 4, 5**). We did not observe any interaction with proximal airway cells **(Figures S9-S11)**. However, the extent of cell interaction differed between materials. In organoids exposed to s/lGO, the materials co-localised with HOPX and MUC1, markers of AECI and ACEII respectively (**Figures 4A-4B**), but this interaction was transient. After 7d exposure, there was very limited cell-material interaction, but rather s/lGO was confined to regions of secretory mucins (MUC1/MUC5AC). We compared the expression of secretory mucins and surfactant in the NM-exposed organoids to the non-injected controls (**Figures 4C-4G**). Interestingly, there was a greater mucin production in response to lGO than sGO exposure (**Figures 4A-4B, 4F)**. In organoids exposed to lGO, we observed a 2.1- and 1.8-fold increase in MUC5AC and MUC1 expression, respectively (**Figure 4F**). In contrast, sGO did not induce mucin hypersecretion in the organoids (**Figures 4A, 4C)**, highlighting the importance of nanosheet lateral dimension. The limited interaction of s/lGO with alveolar cells at 7d also resulted in no alteration in the surfactant homeostasis of the lung organoids (**Figures 4D-4E, 4G)**.

**Figure 4.**
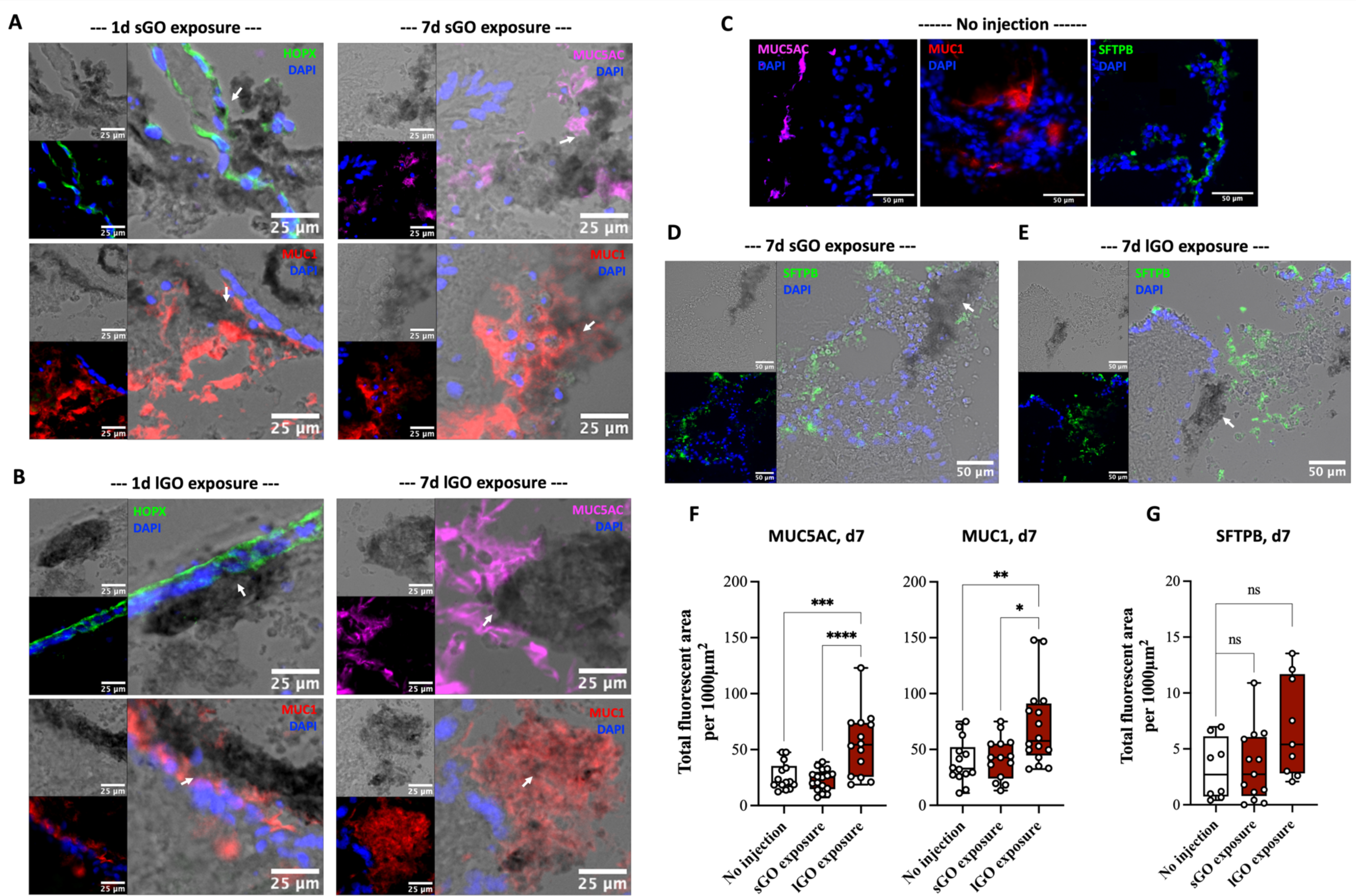
Interaction of sGO and lGO with alveolar cells, secreted mucins and surfactant in lung organoids. **A,** sGO exposure. **B,** lGO exposure. After material exposure, organoids were cryo-sectioned at thickness of 10 µm and immuno-stained for markers of proximal and distal airway epithelial cells. s/lGO dose, 0.2 µg/mm^2^. HOPX (marker for AECI cells); MUC1 (marker for AECII cells when associated to organoid epithelium, or secretory mucin when present in the luminal space); MUC5AC (goblet cells when associated to organoid epithelium, or secretory mucin when present in the luminal space); DAPI (cell nucleus). Arrows indicate material localisation. Scale bar, 25 µm. **C,** Mucin (MUC5AC, MUC1) and surfactant (SFTPB) secretion in the non-injected controls. Scale bar, 50 µm. **D-E,** Surfactant secretion in response to sGO **(D)** and lGO **(E)** exposure for 7d. Scale bar, 50 µm. **F-G,** Semi-quantitative analysis showing the change in mucin (MUC5AC, MUC1) and surfactant (SFTPB) secretion after 7d exposure. Boxplots represent the median, with whiskers indicating minimum and maximum of data (n = 8 to 17 independent images, from 3 experiments). One-way ANOVA \**P <* 0.05, \*\**P <* 0.01, \*\*\**P <* 0.001, \*\*\*\**P <* 0.0001.

In contrast, MWCNT exposure induced a more persistent interaction with AECI/II cells (**Figure 5A**). Regardless of the exposure time, MWCNT remained localised to cells expressing HOPX and MUC1. In these organoids, we did not observe any signs of mucin hypersecretion but rather a 7.8-fold increase in surfactant SFTPB expression, a protein secreted by mature ACEII (**Figures 5B-5C)**. Furthermore, in areas of high material load (indicated by a yellow arrow in **Figure 5A**), cells stained for neither epithelial, proximal nor distal airway cells. Taken together, these findings suggest that nanometric-sized GO has the least impact on HLOs, with MWCNT inducing a more persistent material-cell interaction that alters the surfactant homeostasis of the lung organoid.

**Figure 5.**
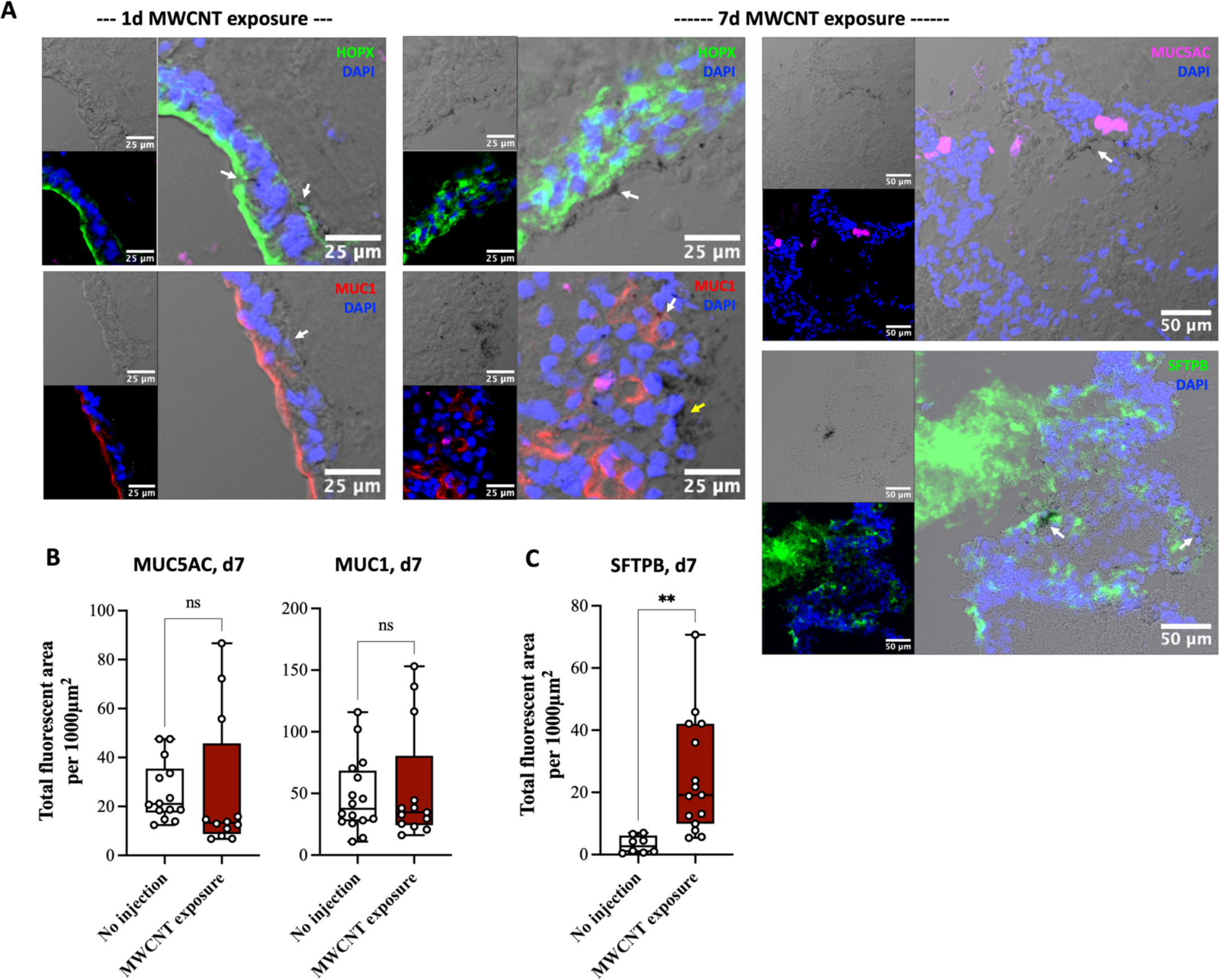
Interaction of MWCNT with alveolar cells, secreted mucins and surfactant in lung organoids. **A,** 1d and 7d exposure to MWCNT. After material exposure, organoids were cryo-sectioned at thickness of 10 µm and immuno-stained for markers of proximal and distal airway epithelial cells. MWCNT dose, 0.01 µg/mm^2^. Organoids exposed to a mid dose of MWCNT were used to assess material-cell interaction due to the detrimental impact of MWCNT on the organoid, when sectioning at high material load. HOPX (marker for AECI cells); MUC1 (marker for AECII cells when associated to organoid epithelium, or secretory mucin when present in the luminal space); MUC5AC (goblet cells when associated to organoid epithelium, or secretory mucin when present in the luminal space); SFTPB (marker for surfactant protein B produced by AECII); DAPI (cell nucleus). Arrows indicate material localisation. **B-C,** Semi-quantitative analysis showing the change in mucin (MUC5AC, MUC1) and surfactant (SFTPB) secretion after 7d exposure. Boxplots represent the median, with whiskers indicating minimum and maximum of data (n = 8 to 16 independent images, from 3 experiments). Unpaired two-tailed Student’s t-test. ns = not significant, \*\**P <* 0.01.

### 3.6 MWCNT induce pulmonary fibrosis in the organoid after 7d exposure

Persistent injury to the alveolar epithelium is known to induce fibrosis, where AECII undergo epithelial-mesenchymal transition (EMT) to repair the injured tissue^[41,42]^. We observed signs indicative of MWCNT-induced pulmonary injury (**Figures 3B, S7B, S8B**), coupled with persistent interaction of MWCNT with the alveolar epithelium (**Figure 5A**), alteration to the lung organoid surfactant homeostasis (**Figure 5C**), and the presence of non-epithelial cells in regions of high MWCNT load (yellow arrow in **Figure 5A**). Such findings promoted an investigation into the biological impact of MWCNT in lung organoids, with particular focus on fibrosis. Unlike Carvalho *et al.*^[39]^, we observed positive staining for markers of mesenchymal cells in HLOs, highlighting the organoid’s potential for modelling pulmonary fibrosis (**Figure 6A**, non-injected controls). To assess if MWCNT interaction with AECI/II leads to fibrosis, we stained for epithelial and mesenchymal markers. In organoids exposed to MWCNT for 7d, but not 1d, there was an increased presence of cells expressing α-SMA (a member of actin family typically expressed by fibroblasts) and vimentin (an intermediate filament and key biomarker of EMT), coupled with a decrease in EpCAM^+^ cells (**Figure 6A-C**). These significant observations at d7 were further confirmed by quantifying the fluorescence area in the images (**Figure 6D**). Together, this indicates the presence of a fibrotic phenotype induced by MWCNT exposure.

**Figure 6.**
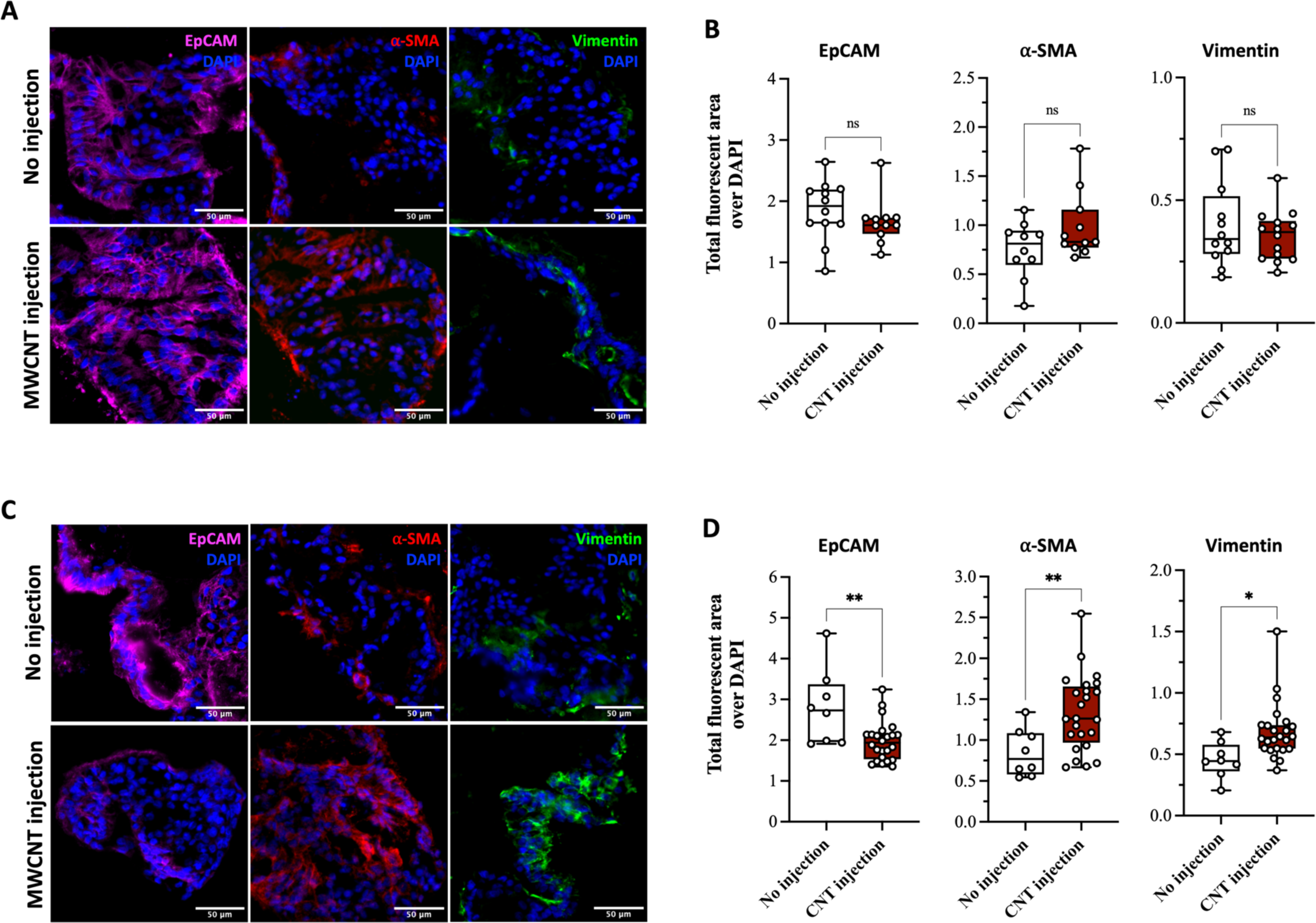
MWCNT-induced fibrosis in lung organoids after 1d and 7d exposure to mid dose MWCNT. **A-B,** 1d exposure. **C-D,** 7d exposure. **A and C**, Immunofluorescence micrographs of lung organoid sections stained for epithelial (EpCAM) and mesenchymal (⍺-SMA, vimentin) markers after MWCNT exposure. **B and D**, Semi-quantitative analysis showing the change in the expression of the indicated epithelial (EpCAM) and mesenchymal (⍺-SMA, vimentin) markers after MWCNT exposure. Boxplots represent the median, with whiskers indicating minimum and maximum of data (n = 8 to 24 independent images, from 3 experiments). Unpaired two-tailed Student’s t-test. ns = not significant, \**P <* 0.05, \*\**P <* 0.01.

## 4.0 Discussion

Advances in stem cell technology and *in vitro* pulmonary tissue culture systems have stimulated the need for more reliable testing approaches, that predict adverse tissue responses and hazard from NM exposures and align with the 3Rs framework^[7]^. Lung organoids, provide a system which supports the co-existence of diverse cell types, an intrinsic dynamic environment created by the presence of functional multi-ciliated and secretory cells, extensive cell-cell, cell-matrix interactions, and tissue-like morphology^[13,14,17,18]^. Though HLOs have been used to better understand various diseases^[12,14,21]^, no studies have attempted to interrogate the potential of functioning, multilineage HLOs as alternative modelling platforms for materials hazard assessment.

In this study, we asked whether hESC-derived lung organoids can model NM-induced pulmonary tissue responses and generate similar outcomes to established *in vivo* models. The HLO development protocol recapitulates both proximal and distal airways in 50d^[18,39]^ (**Figure 1I**). Importantly, it generates functional multiciliated cells that beat (**Movie 1**), mucus-producing goblet cells and surfactant-producing AECII cells (**Figure 1I**), all crucial for creating a true-to-life NM exposure scenario. The genome-wide expression signature of this multilineage organoid also corresponds to adult lung^[18]^. In other protocols, the maturity of HLOs is either unreported or equivalent to foetal lung^[13,14]^.

Microinjection has been previously utilised for presenting pathogens to the apical surface of organoids, as demonstrated in gut organoids infected with *Escherichia coli*^[43]^. The success rate of microinjection is however highly variable between organoids of different sizes, shapes and luminal volumes^[43,44]^. Thus, it was important to develop a reproducible method for determining the organoid size and subsequently the NM dose and volume to microinject. We coupled image analysis of intact organoids with cell counting of matrix-free dissociated organoids (**Figure 2B**). Though the imaging technique only accounts for the 2D cartesian plane (xy), there was an excellent correlation between organoid size and cell number suggesting that the depth (z-plane) of each organoid is proportional to its width (x-plane) and height (y-plane). One group has previously sheared the organoids through mechanical disruption before incubating with desired microparticles and re-embedding the organoids in an extracellular matrix^[45]^. Though it is technically easier than microinjection, this method risks high mechanical injury at the time of incubation, and exposure of both apical and basolateral surfaces. Microinjection thus allowed us to gain access into the lumen of each organoid and dose its apical surface only with a variety of NMs accurately, reproducibly and precisely, without compromising the epithelial barrier or the risk of mechanical injury-induced effects.

We selected two extensively studied carbon-based NMs, GO sheets^[1,4,5,37]^ and MWCNT^[8,24,26,27]^, to validate our HLO exposure model. Thin GO sheets are known to induce minimal-to-transient pulmonary toxicity *in vivo*^[37]^, whilst MWCNT (Mitsui-7) have consistently been associated with adverse pulmonary effects^[27,30,42,46]^. Irrespective of the NM, we observed a transient apoptotic effect in the lung organoids **(Figure S5)**. Interestingly, co-/tri-cultures have been shown to be more resilient to NM exposure than monocultures^[10]^. This suggests that the heterogenous cell population and 3D architecture of the organoid enhances the likelihood of recovery post exposure, and further supports the notion that monocultures might overestimate the response to NMs^[14]^.

Regardless of the primary lateral dimension, sGO and lGO sheets formed agglomerates, trapped in secretory mucins (MUC5AC/MUC1), within the organoid lumen after 7d exposure (**Figures 4A-4B)**. Previous *in vivo* studies have shown that impact of graphene-based materials is size-dependent^[4,5,37]^, and that micrometric GO sheets do not translocate well into lungs and are likely to undergo mucociliary clearance^[37]^. Indeed, we observed increased mucin production in lGO-exposed organoids at d7 (**Figures 4C, 4F)**, suggesting that organoids can tailor the mucin production to reduce damage to the epithelial surface and achieve sufficient mucociliary clearance. Though only a small proportion of lGO sheets reaches the lungs *in vivo*^[37]^, high material reactivity has been previously reported. One factor that has been shown to reduce the adverse biological impact of micrometric materials is protein coating^[1]^. Thus, the overproduction of mucin in response to lGO and its entrapment in secretory mucin may explain the lack of lGO-induced cytotoxicity in the organoids at d7 **(Figure S5B)**. In rodents, lGO but not sGO exposure induces pulmonary granuloma formation, following the recruitment of interstitial macrophages and dendritic cells^[4,37]^. This however resolves within 90d and does not lead to fibrosis^[37]^. We were unable to model the formation of granuloma in response to lGO exposure, as our organoids recapitulate the epithelial component of human lungs. But in agreement with *in vivo* data, organoids exposed to GO did not exhibit any histopathological changes (**Figures 3, S7-S8)**, or remodelling of the extracellular matrix indicative of fibrosis^[37]^. This further highlights the potential of organoids in predicting graphene-based material toxicity.

In contrast to GO, MWCNT exposure caused significant remodelling of lung organoids (**Figures 3B, S7B, S8B)**. CNTs are known to rapidly penetrate the alveoli and interstitial space *in vivo*^[29,41,47]^. Mercer *et al.* showed that MWCNT (80 µg/mouse) penetrates type I epithelial cells within only 1d of exposure^[47]^. Indeed, we observed a persistent interaction of MWCNT with the alveolar epithelium, coupled with limited mucin secretion suggesting a lack of mucociliary clearance (**Figures 5A-5B**). Persistent injury to the alveolar epithelium has previously been shown to lead to lung fibrosis^[37,42]^. Consistent with *in vivo* data, MWCNT exposure for 7d induced EMT, an indirect indicator of fibrinogenic activity of this NM, as demonstrated by an increase in cells expressing mesenchymal markers coupled with a decrease in epithelial cell number (**Figure 6B**).

Pulmonary injury such as fibrosis is also known to alter the homeostasis of the lung^[37,42,48,49]^. *In vivo* studies have shown that inducing lung injury (by intratracheal injection of amiodarone in mice or bleomycin in rats) significantly elevates the surface tension of the lung and/or surfactant SFTPB/C levels^[48],[49]^. Indeed, in MWCNT-treated organoids we observed a significant increase in the expression of SFTPB (**Figures 5A-5C**), a pulmonary surfactant important for decreasing the surface tension along the alveolar epithelium and reducing alveolar collapse^[50]^. Exposure to a pro-fibrotic stimulus like MWCNT may have increased the surface tension of the alveolar epithelium, prompting the surfactant-producing cells in the organoid, AECII, to synthesise more SFTPB. This elevated surfactant level may be an attempt to reduce the surface tension induced by fibrosis and repair the MWCNT-damaged organoid. Alterations in surfactant homeostasis may also explain why organoids exposed to MWCNT lacked a clear lumen, appearing more collapsed (**Figures 3A-4B, S7B-S8B)**. Conversely, exposure to a non-fibrotic stimulus such as GO did not cause any changes in surfactant SFTPB expression (**Figures 4D-4E, 4G)**. This observed consistency between the histopathology and biochemistry endpoints suggests that our model is capable of detecting and responding to pro-fibrotic nanomaterials, such as MWCNT, with relatively high sensitivity and specificity.

## 5.0 Conclusions

In the present study, a HLO exposure model was developed and its suitability as a novel tool for recapitulating pulmonary tissue responses induced by carbon-based NMs was demonstrated. By employing the use of microinjection, we developed a protocol that exposes the apical side of the cells to NMs, emulating human-like lung exposures. We demonstrated that lung organoids can tolerate GO material exposure. Both nanometric and micrometric GO sheets were found to interact transiently with alveolar cells, and were ultimately entrapped in secretory mucin. In contrast, MWCNT induced a more persistent interaction with the alveolar epithelium, that altered the surfactant homeostasis of the organoid and led to a fibrotic phenotype. In such organoids, there was limited mucin secretion, suggesting a lack of mucociliary clearance. Our HLO exposure model, which aligns with the 3Rs framework, has the potential to bridge the toxicological profiles between animals and humans. The model captures the complexity of cell-nanomaterial, cell-cell and cell-matrix interactions induced by exposure to carbon-based NMs. It overcomes simplistic *in vitro* 2D cell models and can potentially provide an ubiquitous and versatile tool to elucidate the mechanisms underlying NM-induced toxicity. With further validation using other carbon/non-carbon NMs, prolonged exposure times, and the incorporation of an immune component, this model could replace current 2D *in vitro* systems, and greatly reduce the need for *in vivo* inhalation studies in the assessment of material pulmonary responses and toxicity.

## Acknowledgments

This work was supported by the GrapheneFlagship project, H2020-FET-GrapheneCore3-#881603. The ICN2 is funded by the CERCA programme / Generalitat de Catalunya and has been supported by the Severo Ochoa Centres of Excellence programme (SEV-2017-0706), and is currently supported by the Severo Ochoa Centres of Excellence programme, Grant CEX2021-001214-S, both funded by MCIN/AEI/10.13039.501100011033. The authors would like to acknowledge Dr Luis M. Arellano and Angeliki Karakasidi for the synthesis and characterisation of GO materials. At the Faculty of Biology, Medicine and Health (University of Manchester), the authors would like to thank the following Core Facility staff for their expert advice and assistant: Dr Peter March and Dr Roger Meadows (Bioimaging Facility), Dr Aleksandr Mironov (Electron Microscopy Facility), Grace Bako and Joel Rigg (Histology Facility), and Dr Gareth Howell (Flow Cytometry Facility). The Bioimaging Facility microscopes and Histology Facility equipment used in this study were purchased with grants from UKRI Biotechnology and Biological Sciences Research Council, The Wellcome Trust and The University of Manchester Strategic Fund. The EM Core Facility (SCR_021147) microscope was purchased with an equipment grant from The Wellcome Trust.

## 6.0 Authors contributions

SV conceived and supervised the work. RI and SV designed the experiments. RI established HLOs, developed the protocol for microinjection, performed all the exposures of HLOs to NMs and analysed all the results. NL provided the GO material and characterisation used in this work. KK supervised preparation and characterisation of GO. Manuscript was written and edited by RI, SV and KK, with the input from NL.

## Abbreviations^1^

^1^ CNT, carbon nanotubes. GO, graphene oxide. HLO, human lung organoids. lGO, large graphene oxide. MWCNT, multi-walled carbon nanotubes. NM, nanomaterial. sGO, small graphene oxide.

