## Supplementary material for "Functioning human lung organoids model pulmonary tissue response from carbon nanomaterial exposures"

**This PDF file includes:**

Figs. S1 to S11

Tables S1 to S3

**Other Materials for this manuscript include the following:**

Movie 1

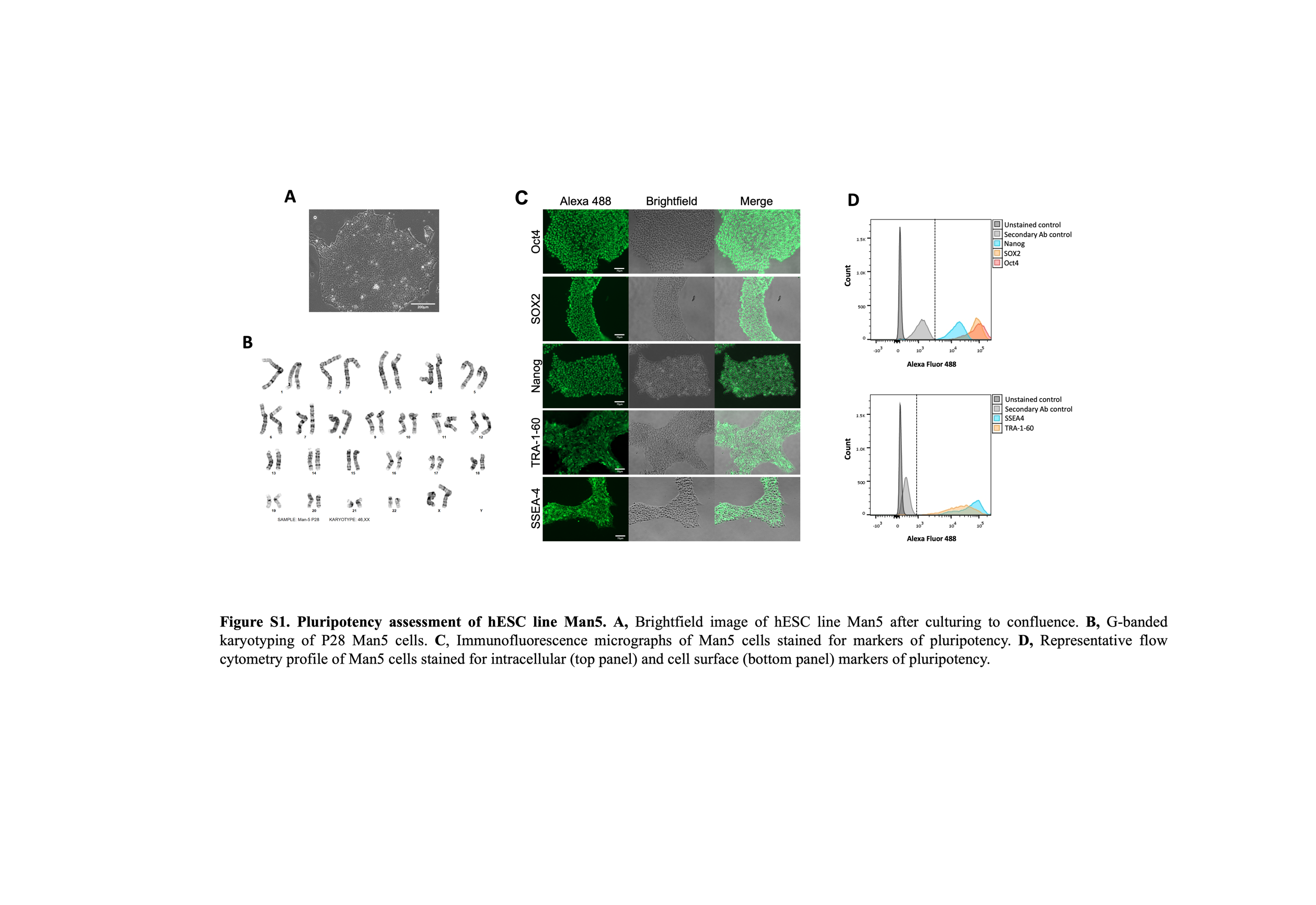

Fig. S1. Pluripotency assessment of hESC line Man5.

A, Brightfield image of hESC line Man5 after culturing to confluence.

B, G-banded karyotyping of P28 Man5 cells.

C, Immunofluorescence micrographs of Man5 cells stained for markers of pluripotency.

D, Representative flow cytometry profile of Man5 cells stained for intracellular (top panel) and cell surface (bottom panel) markers of pluripotency.

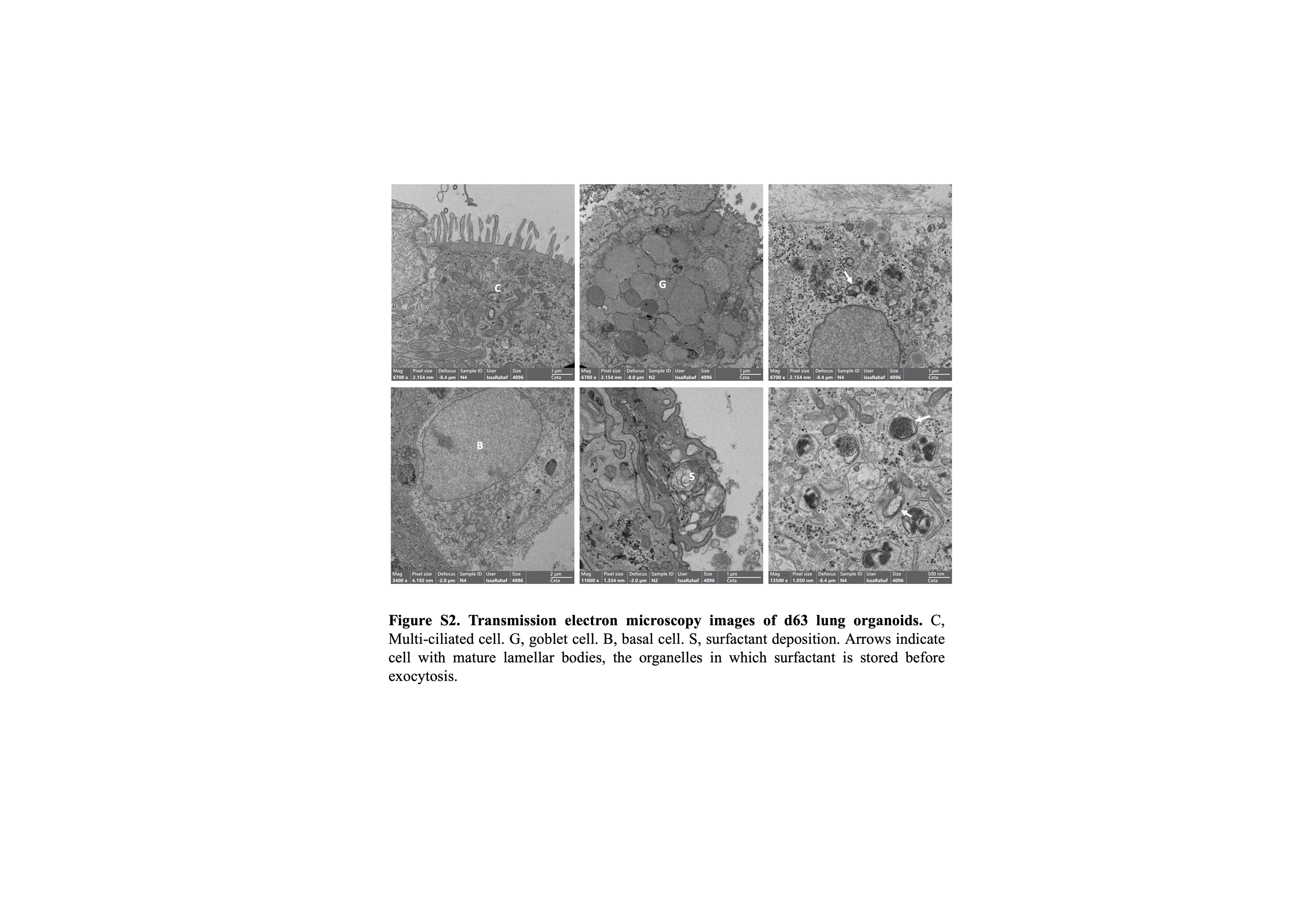

Fig. S2. Transmission electron microscopy images of d63 lung organoids.

Arrows indicate cell with mature lamellar bodies, the organelles in which surfactant is stored before exocytosis. Multi-ciliated cell (C). Goblet cell (G). Basal cell (B). Surfactant deposition (S).

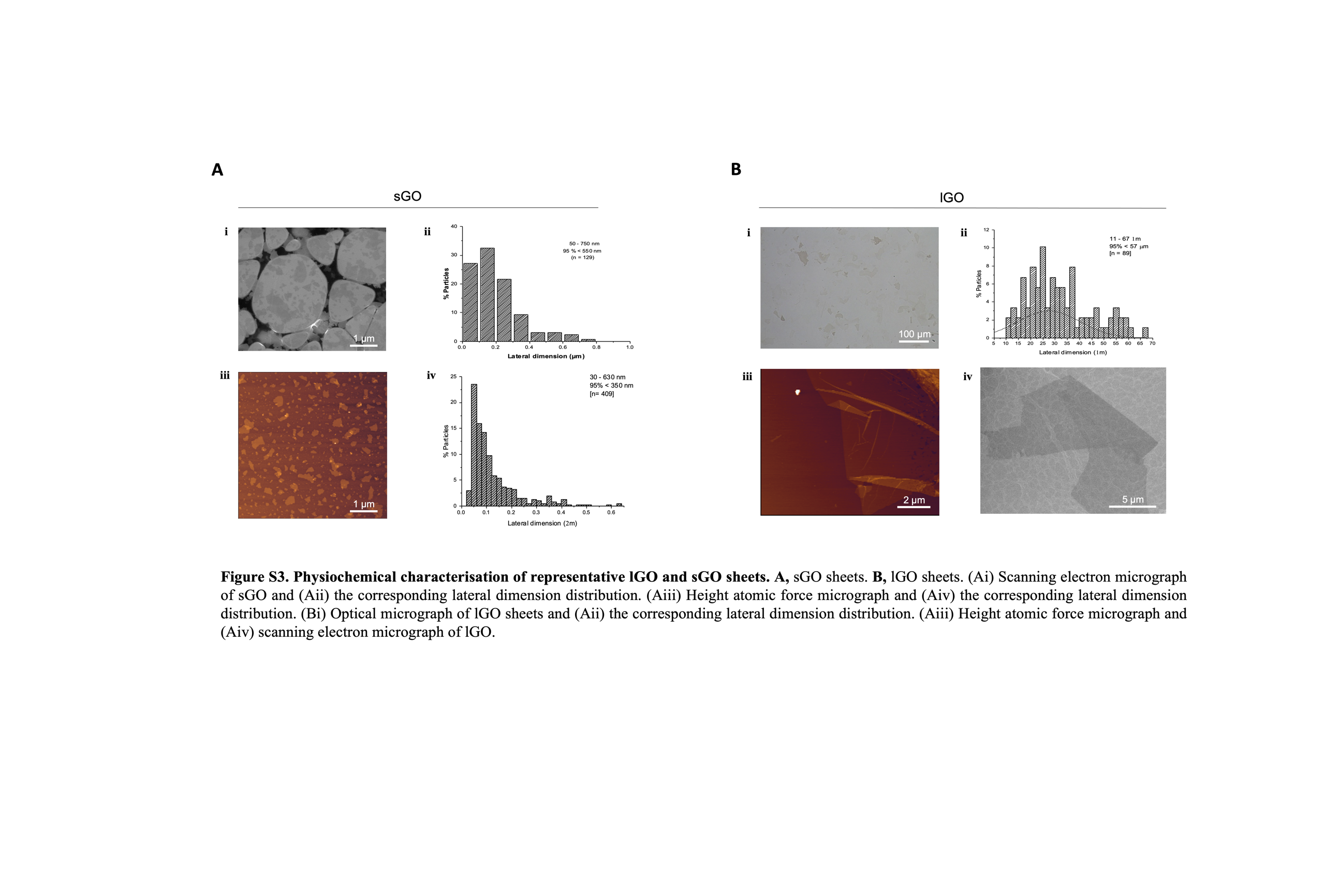

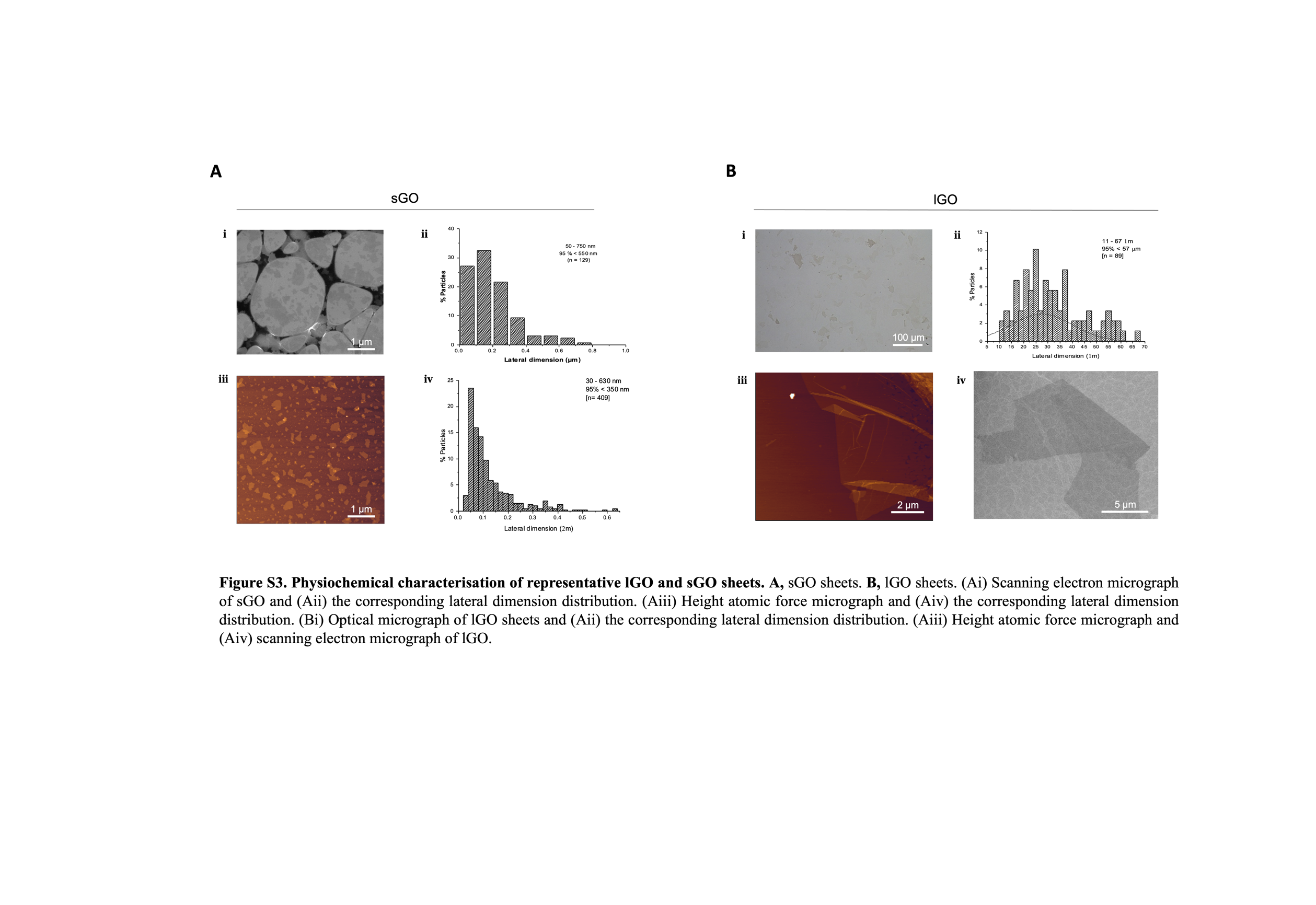

Fig. S3. Physiochemical characterisation of representative lGO and sGO sheets.

A, sGO sheets. B, lGO sheets.

(Ai) Scanning electron micrograph of sGO and (Aii) the corresponding lateral dimension distribution. (Aiii) Height atomic force micrograph and (Aiv) the corresponding lateral dimension distribution.

(Bi) Optical micrograph of lGO sheets and (Bii) the corresponding lateral dimension distribution. (Biii) Height atomic force micrograph and (Biv) scanning electron micrograph of lGO.

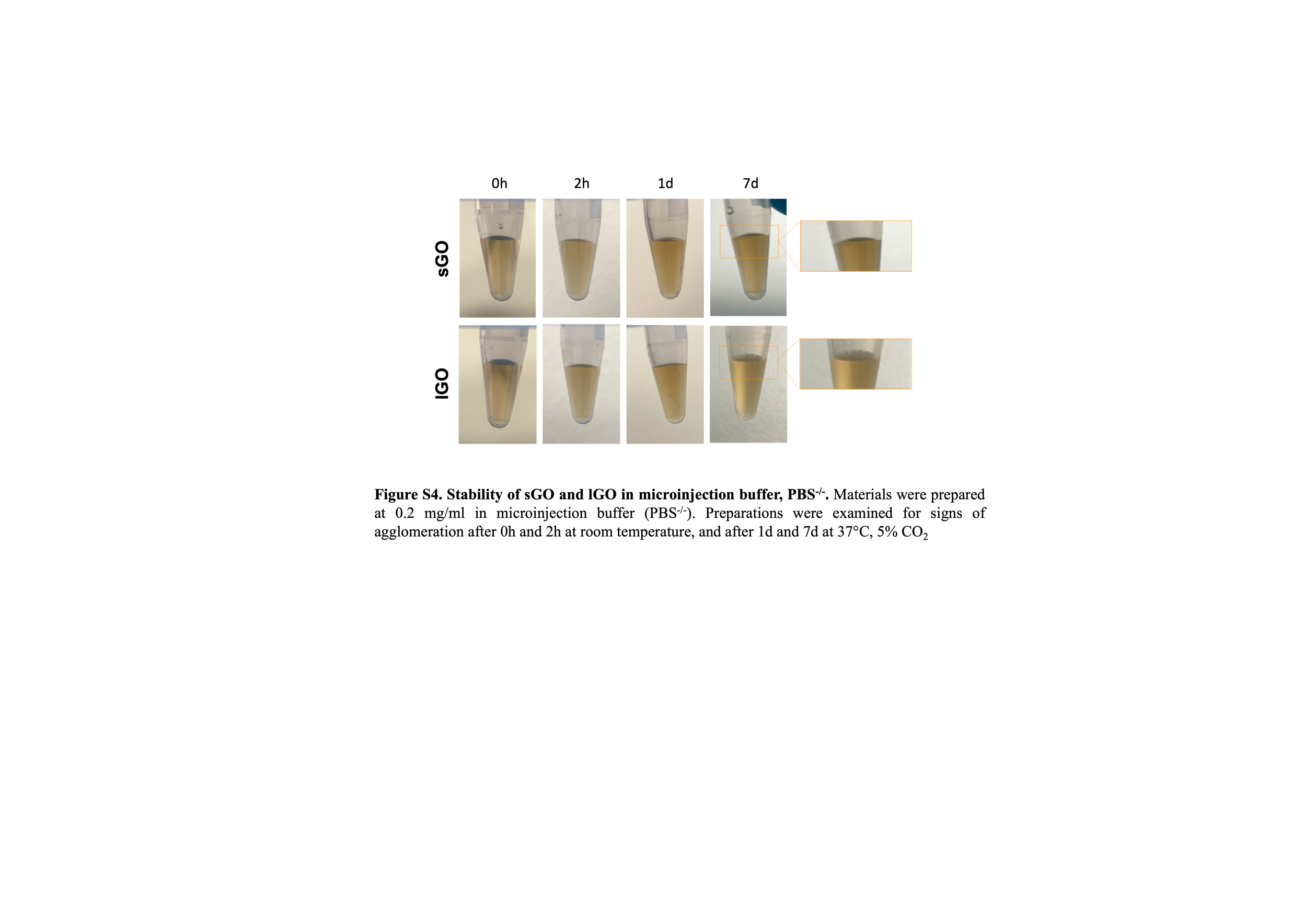

Fig. S4. Stability of sGO and lGO in microinjection buffer, PBS^-/-^.

Materials were prepared at 0.2 mg/ml in microinjection buffer (PBS^-/-^). Preparations were examined for signs of agglomeration after 0h and 2h at room temperature, and after 1d and 7d at 37°C, 5% CO_2_.

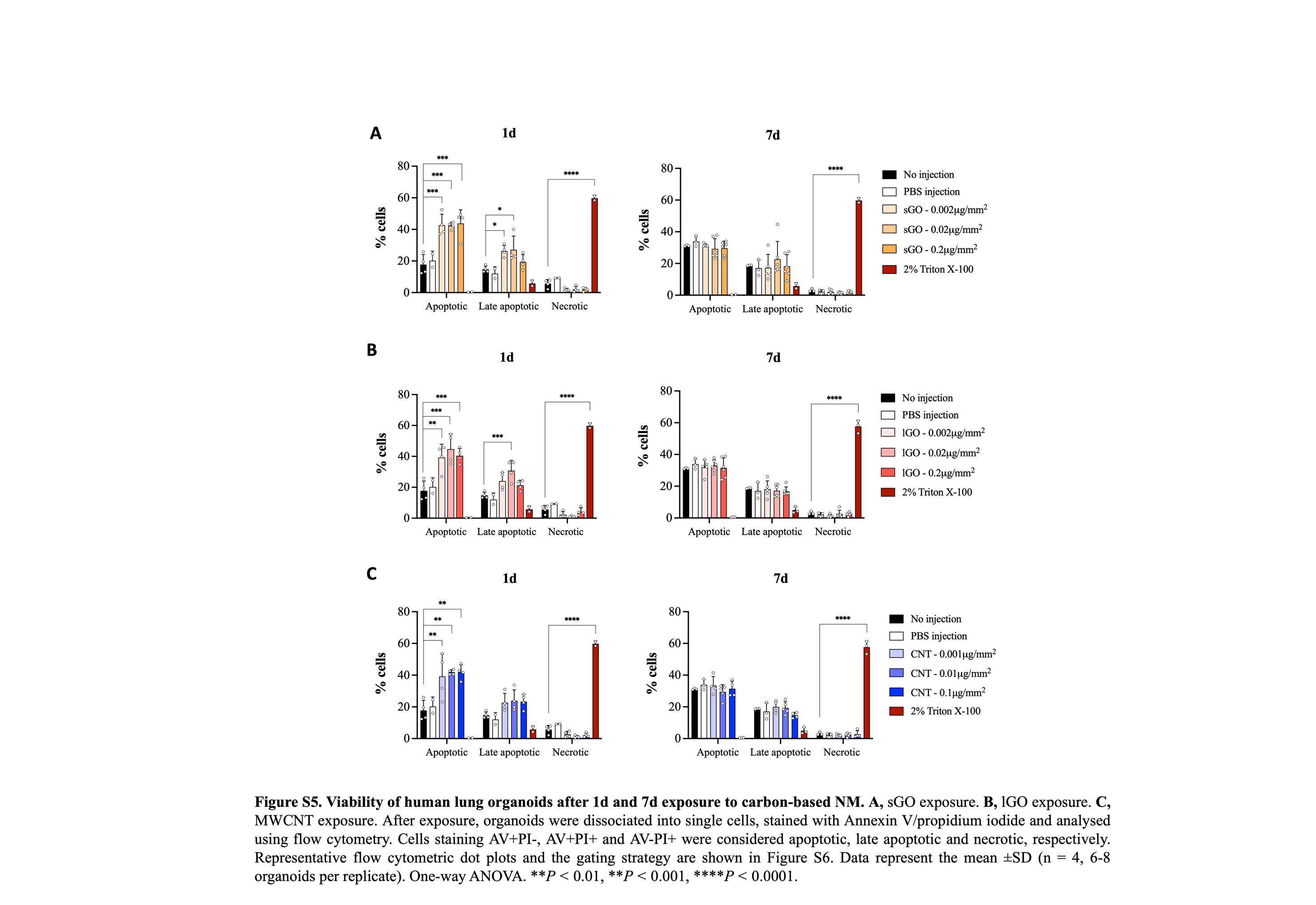

Fig. S5. Viability of human lung organoids after 1d and 7d exposure to carbon-based NM.

A, sGO exposure. B, lGO exposure. C, MWCNT exposure.

After exposure, organoids were dissociated into single cells, stained with Annexin V/propidium iodide and analysed using flow cytometry. Cells staining AV+PI-, AV+PI+ and AV-PI+ were considered apoptotic, late apoptotic and necrotic, respectively. Representative flow cytometric dot plots and the gating strategy are shown in Figure S6. Data represent the mean ±SD (n = 4, 6-8 organoids per replicate). One-way ANOVA. ***P <* 0.01, ***P <* 0.001, *****P <* 0.0001.

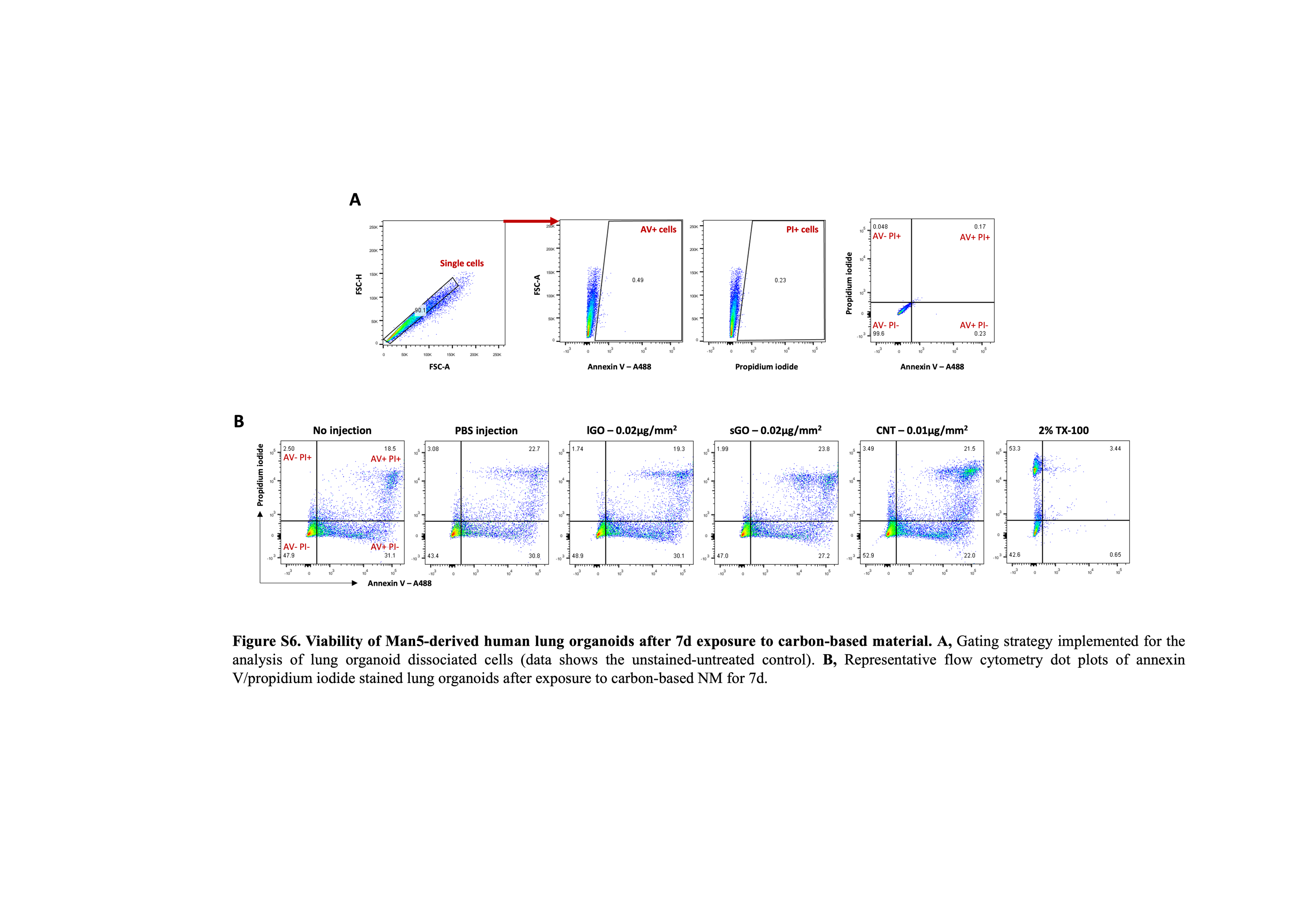

Fig. S6. Viability of Man5-derived human lung organoids after 7d exposure to carbon-based material.

A, Gating strategy implemented for the analysis of lung organoid dissociated cells (data shows the unstained-untreated control).

B, Representative flow cytometry dot plots of annexin V/propidium iodide stained lung organoids after exposure to carbon-based NM for 7d.

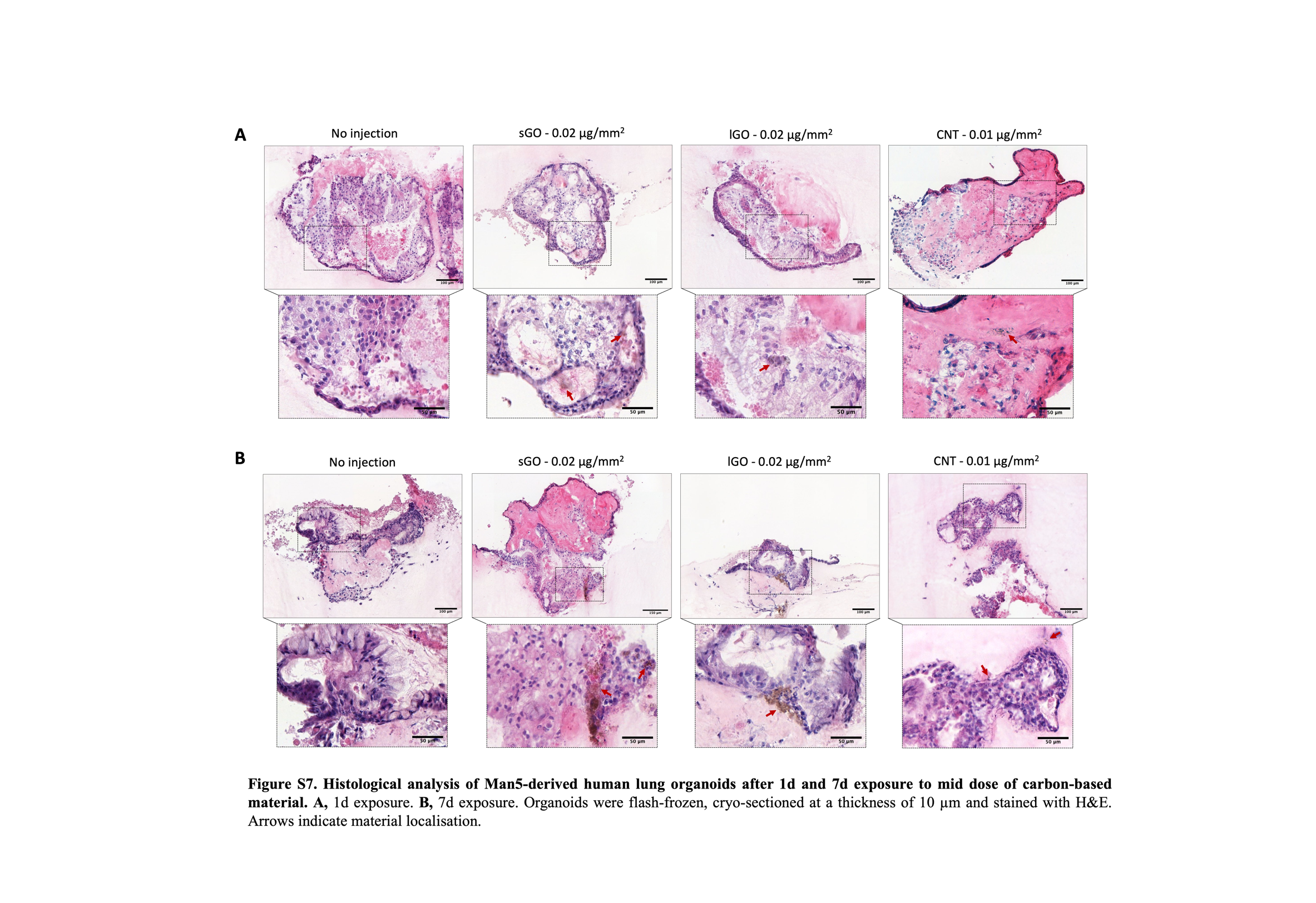

Fig. S7. Histological analysis of Man5-derived human lung organoids after 1d and 7d exposure to mid dose of carbon-based material.

A, 1d exposure. B, 7d exposure.

Organoids were flash-frozen, cryo-sectioned at a thickness of 10 µm and stained with H&E. Arrows indicate material localisation.

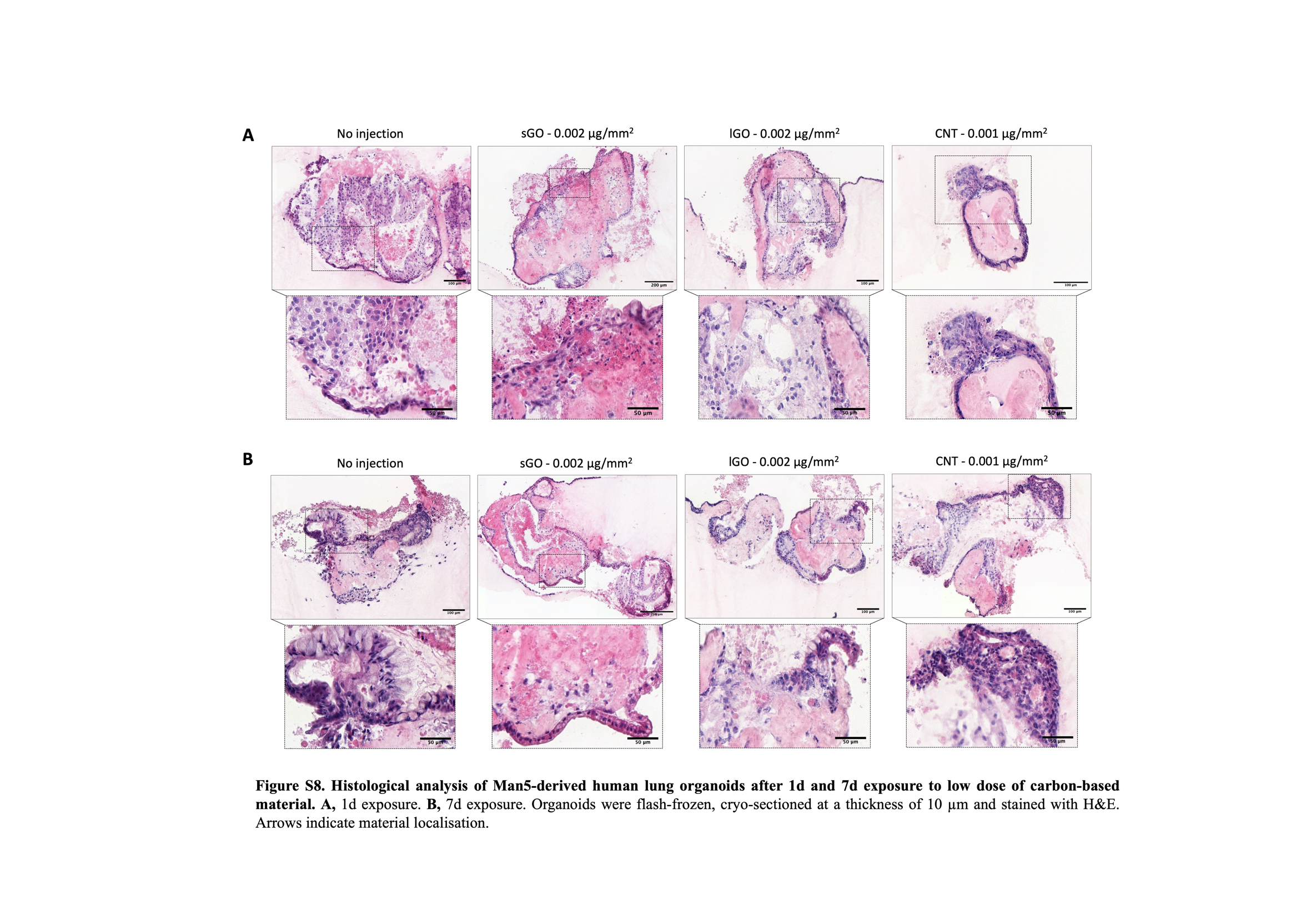

Fig. S8. Histological analysis of Man5-derived human lung organoids after 1d and 7d exposure to low dose of carbon-based material.

A, 1d exposure. B, 7d exposure.

Organoids were flash-frozen, cryo-sectioned at a thickness of 10 µm and stained with H&E. Arrows indicate material localisation.

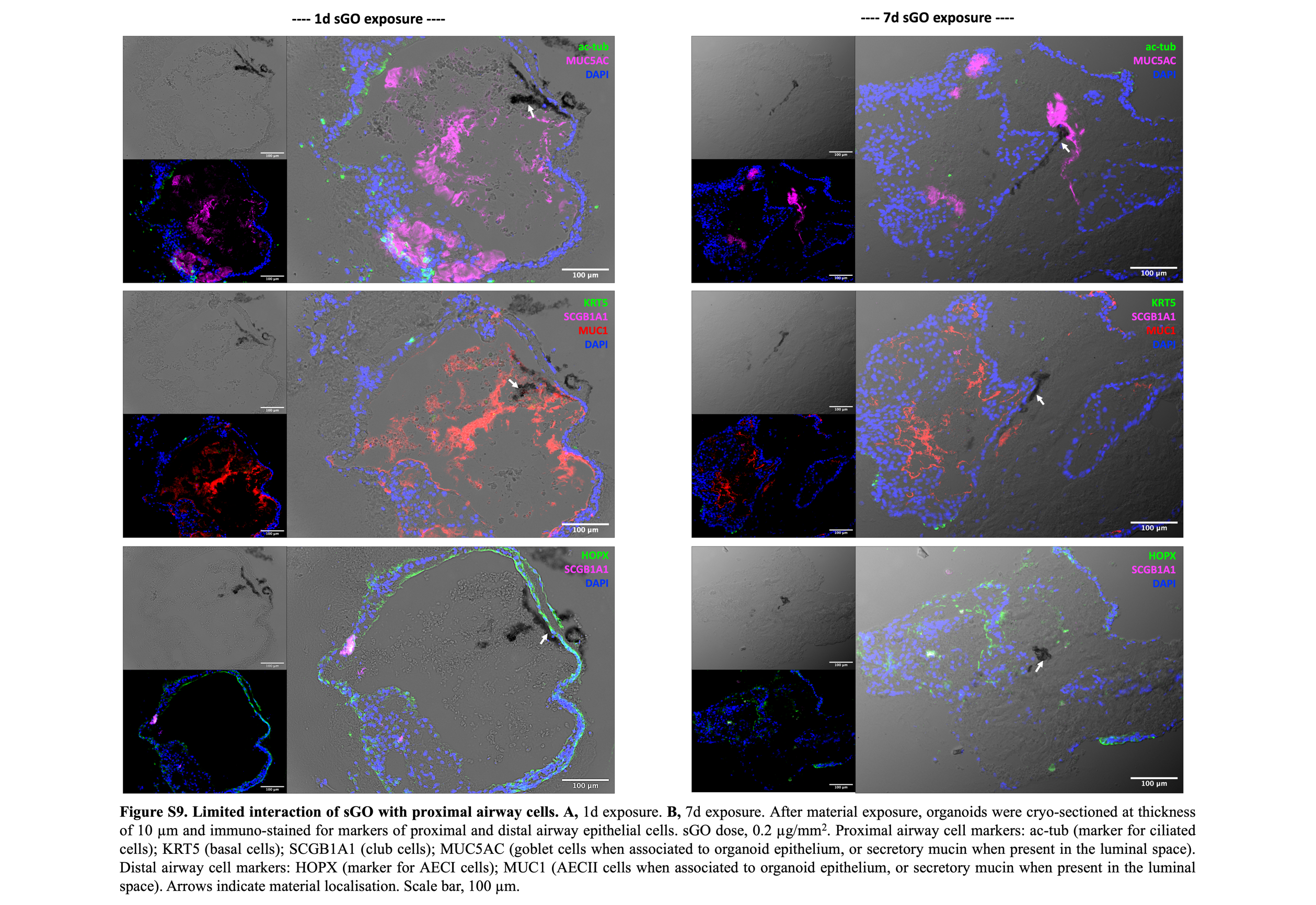

Fig. S9. Limited interaction of sGO with proximal airway cells.

A, 1d exposure. B, 7d exposure.

After material exposure, organoids were cryo-sectioned at thickness of 10 µm and immuno-stained for markers of proximal and distal airway epithelial cells. sGO dose, 0.2 µg/mm^2^. Proximal airway cell markers: ac-tub (marker for ciliated cells); KRT5 (basal cells); SCGB1A1 (club cells); MUC5AC (goblet cells when associated to organoid epithelium, or secretory mucin when present in the luminal space). Distal airway cell markers: HOPX (marker for AECI cells); MUC1 (AECII cells when associated to organoid epithelium, or secretory mucin when present in the luminal space). Arrows indicate material localisation. Scale bar, 100 µm.

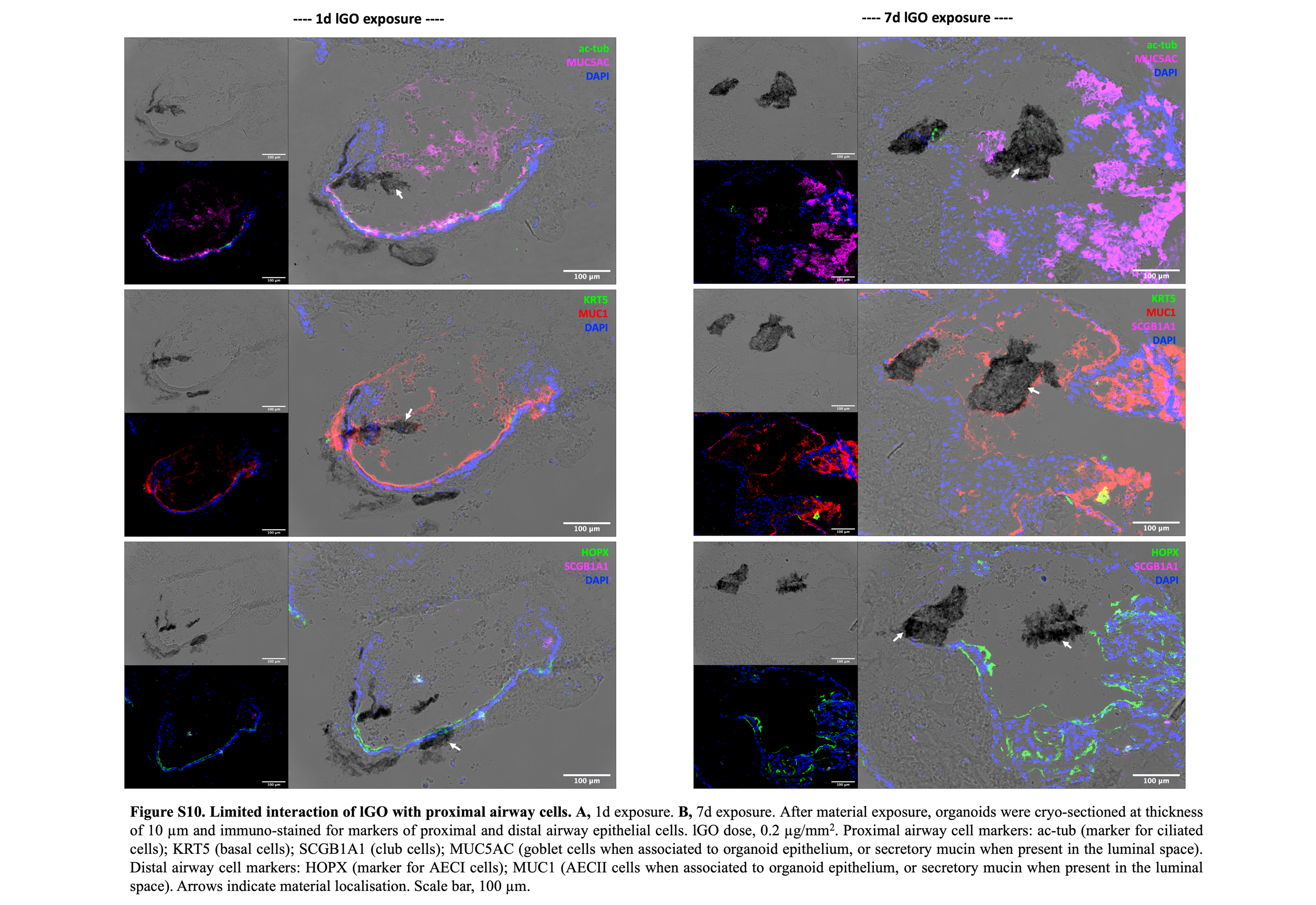

Fig. S10. Limited interaction of lGO with proximal airway cells.

A, 1d exposure. B, 7d exposure.

After material exposure, organoids were cryo-sectioned at thickness of 10 µm and immuno-stained for markers of proximal and distal airway epithelial cells. lGO dose, 0.2 µg/mm^2^. Proximal airway cell markers: ac-tub (marker for ciliated cells); KRT5 (basal cells); SCGB1A1 (club cells); MUC5AC (goblet cells when associated to organoid epithelium, or secretory mucin when present in the luminal space). Distal airway cell markers: HOPX (marker for AECI cells); MUC1 (AECII cells when associated to organoid epithelium, or secretory mucin when present in the luminal space). Arrows indicate material localisation. Scale bar, 100 µm.

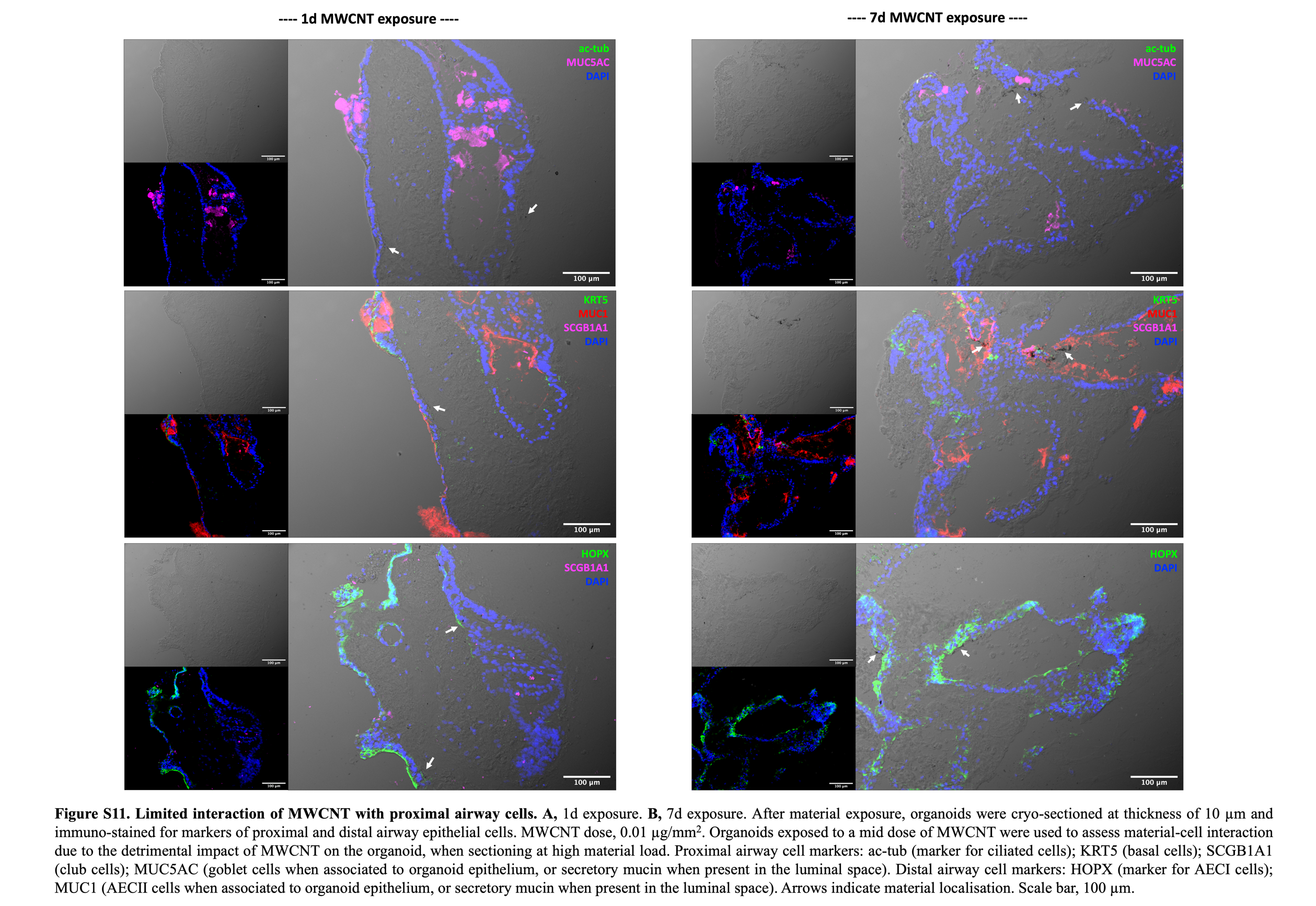

Fig. S11. Limited interaction of MWCNT with proximal airway cells.

A, 1d exposure. B, 7d exposure.

After material exposure, organoids were cryo-sectioned at thickness of 10 µm and immuno-stained for markers of proximal and distal airway epithelial cells. MWCNT dose, 0.01 µg/mm^2^. Organoids exposed to a mid dose of MWCNT were used to assess material-cell interaction due to the detrimental impact of MWCNT on the organoid, when sectioning at high material load. Proximal airway cell markers: ac-tub (marker for ciliated cells); KRT5 (basal cells); SCGB1A1 (club cells); MUC5AC (goblet cells when associated to organoid epithelium, or secretory mucin when present in the luminal space). Distal airway cell markers: HOPX (marker for AECI cells); MUC1 (AECII cells when associated to organoid epithelium, or secretory mucin when present in the luminal space).

**Table S1.** Primary antibodies used for immunostaining.

| ANTIBODY | MANUFACTURER (CATALOGUE NO.) | DILUTION FACTOR |
| --- | --- | --- |
| acTUB | Cell Signalling Technology (#5335) | 1:400 |
| α-SMA | Abcam (#ab5694) | 1:300 |
| EpCAM | Biolegend (#324202) | 1:500 |
| FOXA2 (HNF-3a) | R&D systems (#AF2400) | 1:20 |
| HOPX | Proteintech (#11419-1-AP) | 1:100 |
| KRT5 | Thermo Scientific (#MA5-14473) | 1:200 |
| MUC1 | Thermo Scientific (#MA5-11202) | 1:200 |
| MUC5AC | Thermo Scientific (#MA5-12178) | 1:200 |
| Nanog | Abcam (#ab106465) | 1:100 |
| NKX2.1 | Thermo Scientific (#MA5-13961) | 1:40 (adherent cells);  1:100 (cryosections) |
| Oct.4 | Abcam (#ab19857) | 1:500 |
| SCGB1A1 (CC10) | R&D systems (#MAB4218) | 1:100 |
| SSEA-4 | Abcam (#ab16287) | 1:100 |
| SFTPB (Mature) | Seven Hills (#WRAB-48604) | 1:1000 |
| SFTPC (Mature) | Seven Hills (#WRAB-76694) | 1:500 |
| SOX2 | BD Bioscience (#561469) | 1:50 |
| SOX9 | Sigma Aldrich (#AB5535) | 1:200 (adherent cells);  1:300 (cryosections) |
| TRA-1-60 | Abcam (#ab16288) | 1:100 |
| Vimentin | Abcam (#ab24525) | 1:400 |

**Table S2.** Secondary antibodies used for immunostaining.

| ANTIBODY | MANUFACTURER (CATALOGUE NO.) | DILUTION FACTOR |
| --- | --- | --- |
| Donkey anti-goat 488 | Thermo Scientific (#A11055) | 1:500 |
| Donkey anti-rabbit 488 | Thermo Scientific (#A21206) | 1:500 (adherent cells);  1:200 (flow cytometry);  1:400 (cryosections) |
| Donkey anti-mouse 647 | Thermo Scientific (#A31571) | 1:400 |
| Goat anti-chicken 488 | Thermo Scientific (#A11039) | 1:400 |
| Goat anti-mouse 488 | Thermo Scientific (#A11001) | 1:500 (adherent cells);  1:200 (flow cytometry);  1:400 (cryosections) |
| Goat anti-rabbit 488 | Thermo Scientific (#A11034) | 1:400 |
| Goat anti-hamster 568 | Abcam (#ab175716) | 1:400 |
| Goat anti-rabbit 594 | Thermo Scientific (#A11012) | 1:400 |
| Goat anti-mouse 647 | Thermo Scientific (#A21236) | 1:400 |
| Goat anti-rat 647 | Thermo Scientific (#A21247) | 1:400 |

**Table S3.** Antibodies used for flow cytometry.

| ANTIBODY | CONJUGATE | MANUFACTURER (CATALOGUE NO.) | DILUTION FACTOR |
| --- | --- | --- | --- |
| cKIT | PE | Biolegend (#313204) | 1µl/2x10^5^ cells |
| CXCR4 | APC | Biolegend (#306510) | 1µl/2x10^5^ cells |
| EpCAM | AF488 | Biolegend (#324210) | 1µl/2x10^5^ cells |
| MUC1 | APC | Biolegend (#337004) | 1µl/2x10^5^ cells |
| Nanog | Uncojugated | Thermo Scientific (#MA1-017) | 1:200 |
| Oct.4 | Unconjugated | Abcam (#ab19857) | 1:500 |
| PDNP | PE | Biolegend (#355608) | 1µl/2x10^5^ cells |
| SSEA-4 | Unconjugated | Abcam (#ab16287) | 1:500 |
| SOX2 | Unconjugated | BD Bioscience (#561469) | 1:200 |
| TRA-1-60 | Unconjugated | Abcam (#ab16288) | 1:500 |
